# Models trained with noisy genomes extend bacterial phenotype prediction into deep time

**DOI:** 10.64898/2026.06.29.735159

**Authors:** Anzhelika Koldaeva, Olga Bagrova, Joshua A. M. Mitchell, Philip Hugenholtz, Anja Spang, Ben Woodcroft, Tom A. Williams, Gergely Szöllősi

## Abstract

Predicting phenotype from genotype in extant organisms is increasingly tractable through the accumulation of genome sequences and the development of machine-learning algorithms. Here we show that machine learning can be applied to reconstructed ancestral gene content, extending these predictions into the past. We trained models on a diverse set of bacterial phenotypes and found that introducing noise into gene content profiles allows predictions to generalize over larger evolutionary distances. For phenotypes with signal spread across many genes - such as metabolic oxygen use, cell envelope architecture and optimal growth temperature - noise augmentation extends resolution back to the root of the bacterial domain, while for other phenotypes - including GC content and sporulation - the range remains more limited. We therefore conclude that the last bacterial common ancestor (LBCA) was likely an anaerobic, double-membraned, and moderately thermophilic bacterium (46-75°C). Moreover, this work provides a general approach for learning about the genomic basis of phenotypes and drawing inferences about their early evolution.

## I. Introduction

Understanding how genome features give rise to organismal phenotypes is a central question in evolutionary biology [1], [2], [3]. In microbes, phenotypes such as metabolism, oxygen use, cell envelope architecture, and temperature preference define ecological niches and shape evolutionary success in a given environment.

Prediction of microbial phenotypes from genomic data has traditionally relied on comparative genomics: identifying specific genes and pathways through curated functional annotations linked to a target phenotype [4], [5]. These approaches work well for traits whose genetic basis is already known, but lose resolution when a phenotype is encoded by many genes distributed across the genome [6], [7], or when functional annotations are incomplete or biased by the absence of close homologs in model organisms [8], [9].

The accumulation of microbial genome sequences has enabled a shift toward data-driven approaches. Machine-learning (ML) models learn predictive patterns directly from genome-wide features, integrating information across thousands of genes and capturing non-linear and partially redundant associations that are difficult to identify by hand [10], [11]. ML has been applied to a broad range of microbial phenotypes, including antibiotic resistance [12], [13], metabolic traits [14], [15], and environmental adaptations [16].

Phenotypes are encoded in genomes in many ways: through the presence - or sequence - of individual genes, combinations of genes, gene dosage, and broader regulatory architecture. How this information is encoded and distributed across the genome remains an open question. From an ML perspective, prediction that generalizes requires genomic patterns that remain informative beyond closely related taxa and across deep evolutionary distances, rather than lineage-specific shortcuts that inflate accuracy on near relatives [10], [17]. Whether a phenotype’s signal is concentrated in a few key genes or distributed redundantly across many, how much of the genome is needed to recover it, and how robust it is to gene loss are therefore not only questions of model interpretation: they determine whether the signal persists across lineages and how far back in evolutionary time a phenotype can still be read from genomic data.

Robust phenotype predictors also enable inference beyond extant organisms, in particular the prediction of phenotypes for ancestral genomes [18], [14]. Such inferences can reconstruct how key biological traits have evolved through time and link phenotypic change to environmental and biological conditions across Earth history. In this setting, models are trained on present-day genomes and applied to reconstructed ancestral ones. However, strong performance on extant data does not guarantee reliable ancestral inference, for two reasons. First, predictive power decreases over deep evolutionary distances, both because models generalize poorly and because the way a phenotype is encoded can itself shift over time [19], [20]. Second, reconstructed ancestral genomes are inherently incomplete and noisy, which can obscure even persistent phenotype signals [21], [22]. Reliable ancestral prediction therefore requires understanding how phenotype signal is distributed across the genome and how robust it is to gene loss and noise.

In this work we focus on five microbial phenotypes that span distinct biological mechanisms and levels of predictive difficulty: oxygen use, cell envelope structure, sporulation ability, optimal growth temperature (OGT), and genomic GC content. Together they provide a diverse testbed for studying genome-based phenotype prediction across evolutionary scales. We represent each genome by gene family copy numbers, using COGs assigned by eggNOG-mapper [23], [24] because they map robustly across prokaryotic genomes. This representation provides a compact summary of functional gene content and matches the output of ancestral-genome reconstruction methods based on phylogenetic reconciliation [21], [25], [26]. Training predictors directly on copy-number profiles therefore makes them applicable to reconstructed ancestral genomes, enabling phenotype inference beyond extant taxa and across deep timescales.

To investigate how phenotypes can be predicted across deep evolutionary distances, and where in the genome they are encoded, we have built the pipeline in Fig. 1. We start with extant phenotype-annotated genomes, represented as gene family copy numbers, and systematically evaluate predictive performance under taxonomy-aware train–test splits that progressively increase the evolutionary distance between training and test sets. We then examine how phenotype signals are distributed across the genome using feature subsampling and feature-importance analyses. Next, we train noise-robust models by augmenting training data with noisy genome replicates. Finally, we apply these models to incomplete, noisy genomes, including reconstructed ancestral ones, to infer ancestral phenotypes across the bacterial tree.

**Figure 1:**
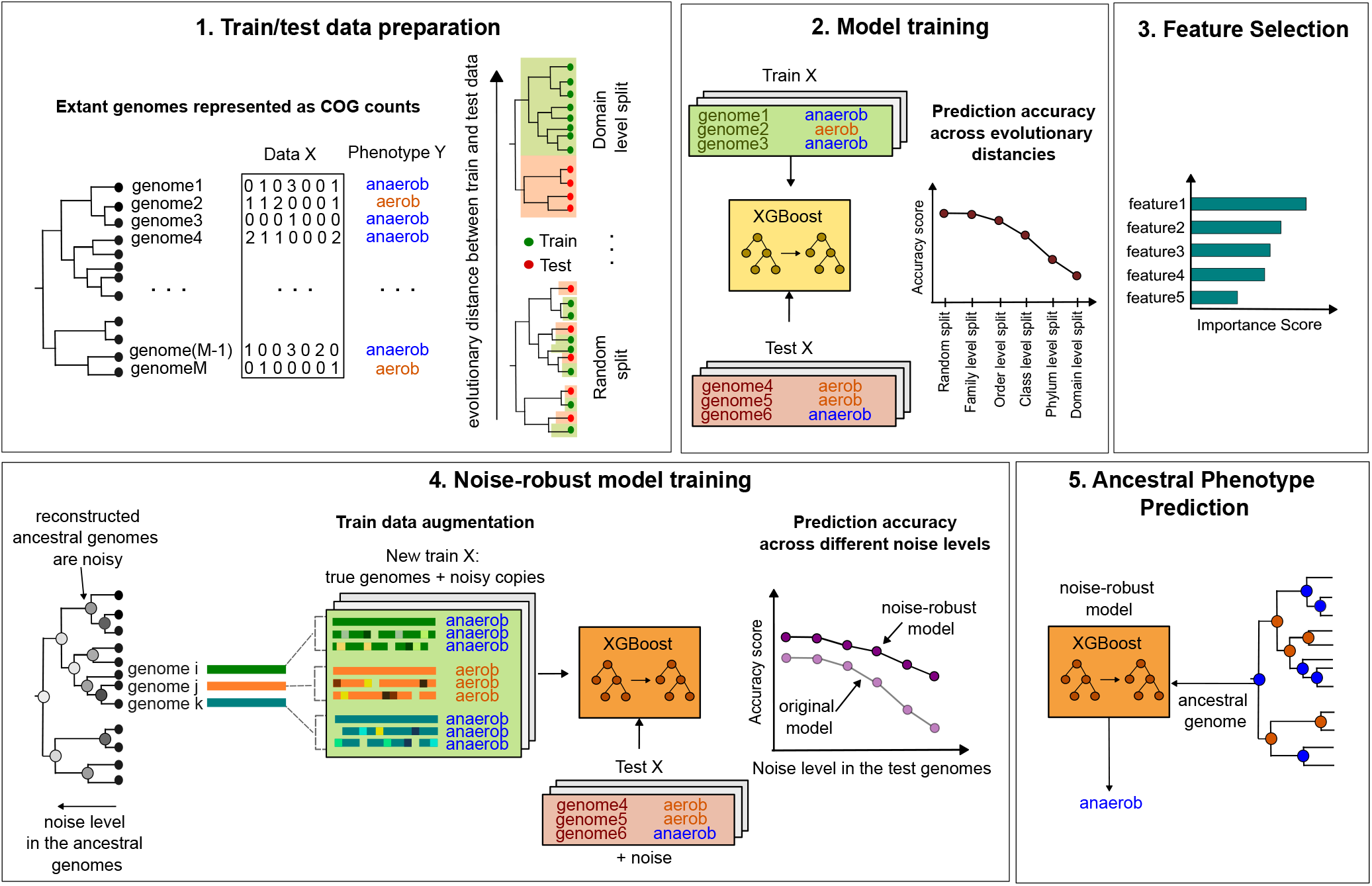
Schematic overview of the pipeline for phenotype signal exploration across evolutionary distances, feature selection, and ancestral phenotype prediction.

## II. Results

### A. Data preparation and phenotype prediction across evolutionary distance

We use five datasets to train and evaluate our predictors. Each dataset is a count matrix *X* whose rows are genomes and whose columns are homologous gene families, defined as Clusters of Orthologous Groups (COGs) [5] and assigned to Genome Taxonomy Database (GTDB) genomes [27] using eggNOG-mapper [23]. Phenotype labels *Y* are obtained by integrating annotations from multiple external sources, described separately for each phenotype below. Projecting the count data into two dimensions with t-distributed stochastic neighbor embedding (t-SNE) [28], we recover known phylogenetic structure, with clusters corresponding to major taxonomic groups, see Fig. 1 in Supplementary Material. Phenotype labels often separate cleanly in this embedding, suggesting that phenotype signal is reflected in genome-wide feature patterns and is at least partially separable in the original feature space [29]. The distribution of samples and labels across phyla, however, is highly imbalanced, motivating phylogenetically informed train–test splits.

Models trained on closely related genomes can exploit lineage-specific shortcuts that inflate performance on randomly sampled test sets. To assess generalization across evolutionary distances, we perform taxonomy-aware train–test splits at five taxonomic levels (family, order, class, phylum, and domain), generating 30 independent splits per level and phenotype.Given the substantial class imbalance, we use the Matthews correlation coefficient (MCC) as the primary performance metric, complemented by balanced accuracy, recall, F1 score, and the Expected Calibration Error (ECE) to assess the calibration of predicted probabilities. Since some phenotypes show high within-group homogeneity, see Supplementary Section 2; Figs. 2, and 3, a phylogeny-only predictor could perform reasonably well by exploiting shared ancestry rather than genomic content. To distinguish these cases, we compared XGBoost predictions against a phylogeny-based predictions that infers each test genome’s phenotype from its position on the GTDB tree alone, see Methods, Section IV-B), using identical train–test splits and evaluation metrics. Results for each phenotype are summarized below.

**Figure 2:**
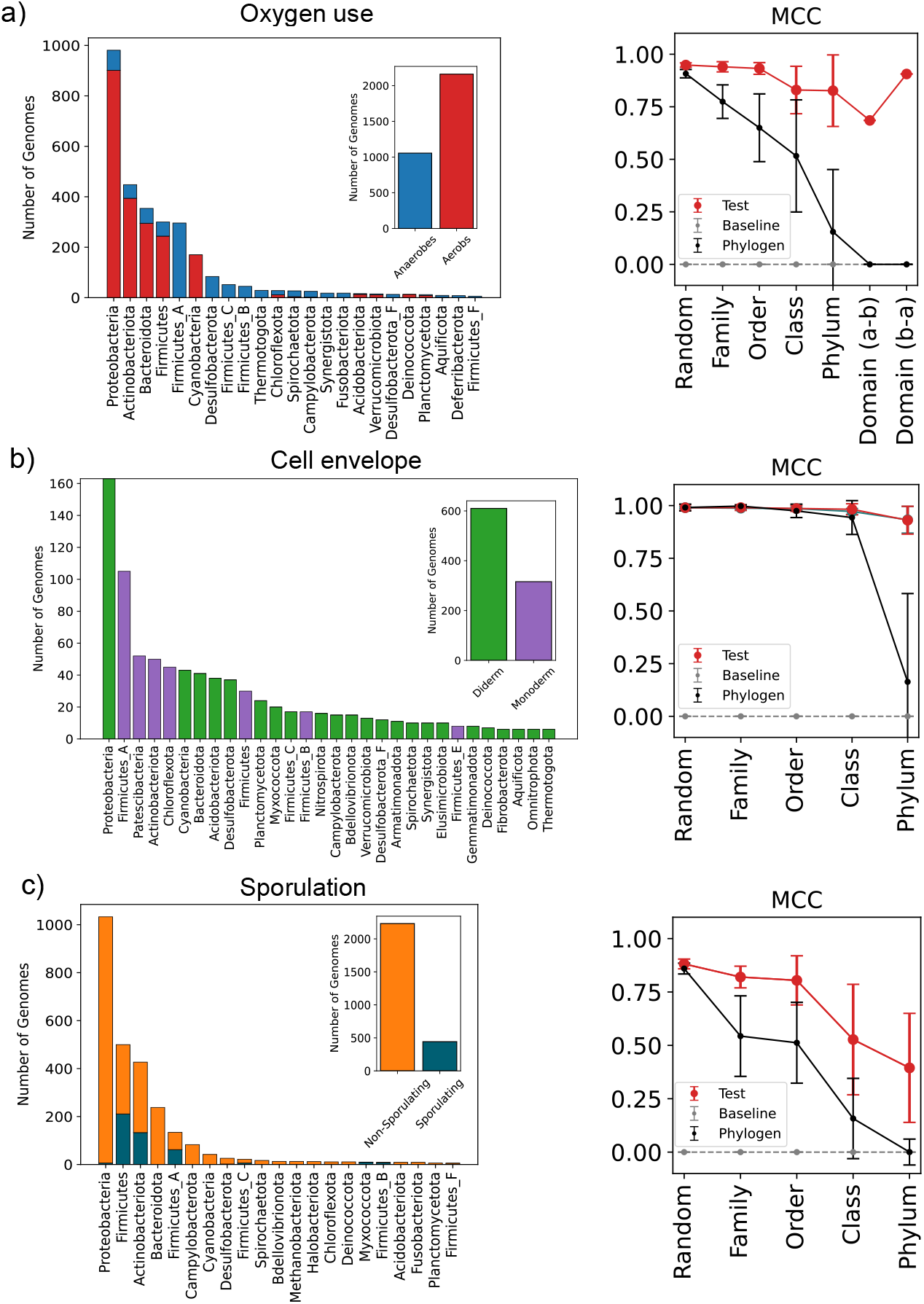
Distribution of genome counts per phylum for different phenotype datasets and MCC scores of phenotype predictors across evolutionary distance (black: phylogeny based prediction; red: model performance on test data; light grey: baseline performance of random guessing). a) Oxygen use phenotype. b) Cell envelope phenotype. c) Sporulation phenotype.

**Figure 3:**
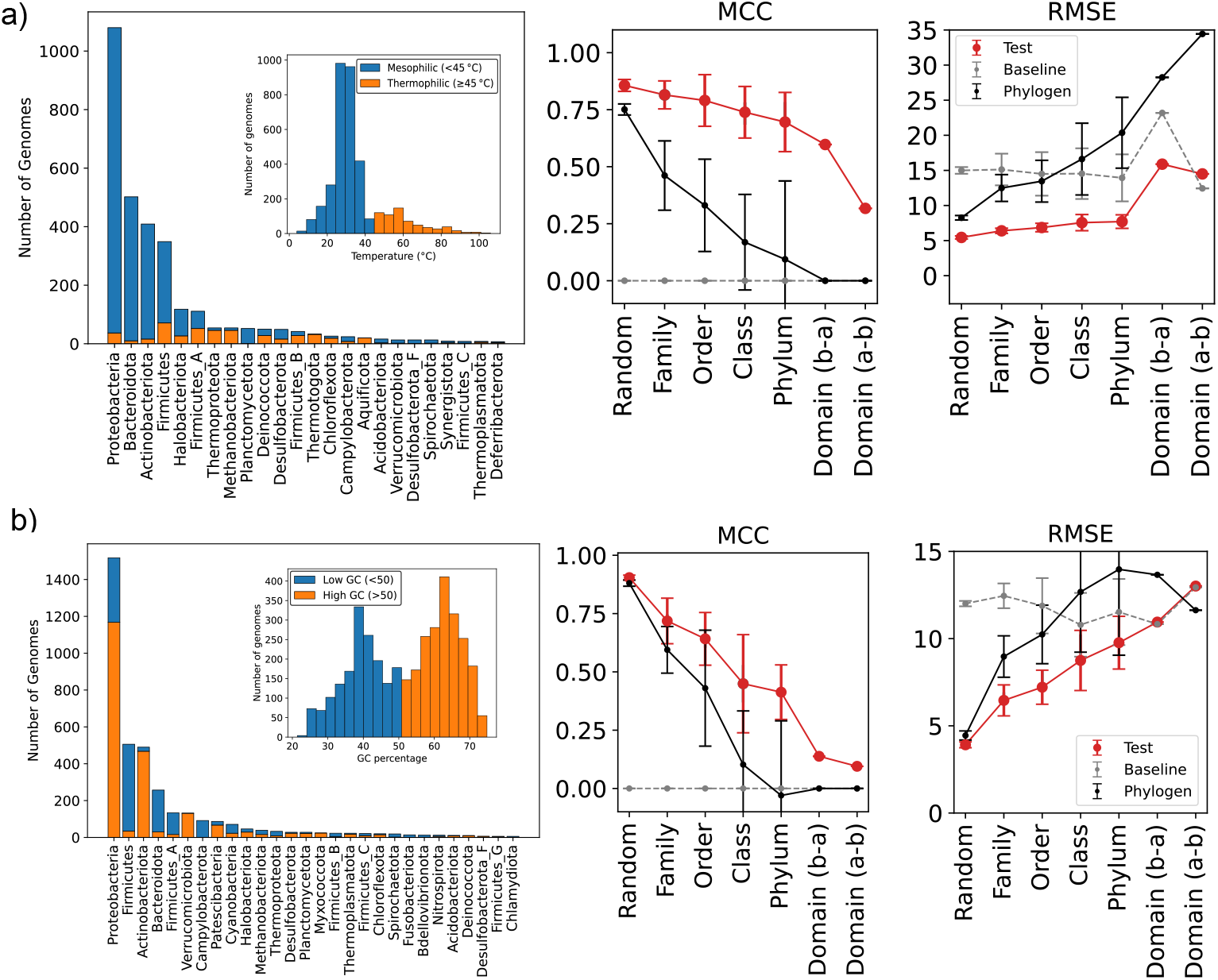
Distribution of genome counts per phylum for different phenotype datasets, MCC and RMSE scores of phenotype predictors across evolutionary distance (black: phylogeny based prediction; red: model performance on test data; light grey: baseline performance of random guessing). a) Optimal growth temperature. b) GC content.

#### 1) Oxygen use

We here define oxygen use as the ability of organisms to grow in the presence of molecular oxygen. It has recently been predicted from genomic data based on a machine learning classifiers using genomic features at multiple levels of resolution, including Pfam protein families [30], [16], annotation-free amino-acid triplet representations [31], and COG-based genome profiles [14]. Here, we use GTDB genomes annotated with oxygen-use information from the BacDive database [32] and focus on the binary classification of aerobic and anaerobic organisms. We extend the framework of [14] by evaluating prediction performance across large evolutionary distances.

Aerobes substantially outnumber anaerobes in our dataset, see Fig. 2(a), left, yet XGBoost maintains strong predictive performance with well-calibrated confidence estimates even under domain-level holdout splits, see Fig. 2(a), right; Supplementary Table 1, where phylogeny-based predictions show a rapid decline in accuracy. This contrast suggests that the learned signal reflects functional genomic properties of oxygen use rather than mere phylogenetic proximity, generalizing across deep evolutionary distances.

#### 2) Cell envelope

The cell envelope is a deeply conserved physiological trait used to broadly classify bacteria as monoderms (single membrane, thick peptidoglycan) or diderms (inner and outer membranes, thin peptidoglycan). This classification broadly, though imperfectly, overlaps with Gram-positive and Gram-negative staining [33], [34].

We formulate a binary classification task distinguishing between monoderm and diderm organisms. Genomes in the 1007-taxon dataset from [14] were classified as monoderm or diderm based on a literature search [35]. The dataset is highly biased toward diderm organisms, with most phyla being exclusively monoderm or diderm, see Fig. 2(b), left. Despite substantial class imbalance, cell envelope phenotype remains highly predictable under deep taxonomic holdout splits, reflecting broadly conserved genomic signatures across bacterial diversity. Model calibration was also strong across splits, suggesting that predicted probabilities are reliable and not merely a consequence of class imbalance, see Fig. 2(b), right. Notably, phylogeny-based predictions also perform well up to the phylum level, suggesting that monoderm/diderm identity largely follows deep phylogenetic structure with limited within-phylum variation.

#### 3) Sporulation

Sporulation is a survival strategy in which some bacteria produce dormant, highly resistant spores in response to unfavorable conditions [36], [37].

We used sporulation annotations from [38] and formulated a binary classification task distinguishing sporulating from non-sporulating organisms. In contrast to oxygen use and cell envelope structure, sporulation is phylogenetically restricted, with sporulating taxa concentrated within a limited number of phyla and most samples annotated as non-sporulating, see Fig. 2(c), left. Accordingly, predictive performance declines sharply at class and phylum level holdouts, see Fig. 2(c), right; Table 3), despite models remaining well calibrated across splits, suggesting the model captures lineage-specific genomic signatures rather than universally conserved sporulation signals, which is consistent with the known restriction of sporulation to specific bacterial lineages [37]. Notably, cell envelope biosynthesis and sporulation are evolutionarily linked, with multiple independent co-losses of both traits observed across the class Bacilli, suggesting shared regulatory and biosynthetic dependencies between the two phenotypes.

#### 4) Optimal growth temperature (OGT)

Unlike the previous binary phenotypes, optimal growth temperature (OGT) is a continuous trait reflecting organismal adaptation to environmental temperatures that impose constraints on microbial metabolism, protein stability, and cellular organization [39]. Predicting OGT from genomic data has been widely explored using machine learning approaches that leverage features such as amino acid composition [40], and protein family profiles [16], [41].

Growth temperature optima and ranges were sourced from the BacDive [32] and TEMPURA [42] databases. Annotations from primary literature were used when multiple sources were available and only species with at least 3 recorded growth temperatures were included.

To predict OGT, we used a two-step XGBoost approach in which genomes were first classified as mesophiles or thermophiles, and the final temperature estimate was calculated as a probability-weighted combination of predictions from the corresponding regressors, see Eq. 2. The binary classifier remains well calibrated across splits, and the performance of both the classifier and the final regression model is summarized in Fig. 3(a), right and Table 4. Prediction accuracy decreases gradually with increasing evolutionary distance between the training and test sets, indicating reduced transferability across deep phylogenetic splits.

#### 5) GC content

Finally, we examined GC content, a genomic property reflecting underlying mutational biases, genome stability, and long-term evolutionary processes [43]. GC content has frequently been used as an informative feature in machine learning models for predicting microbial traits such as optimal growth temperature, habitat, and taxonomy [44], [45].

In this study, we use GC annotations from the prokaryotic trait database of [38] and adopt a similar two-step approach as for OGT predictions to predict continuous GC content values, see Eq. 3. The results of the binary classifier and the final regression model are summarized in Fig. 3(b), right and Table 5. Both predictive performance and model calibration decrease with increasing evolutionary distance, indicating that the model largely relies on lineage-specific correlations with GC content and is less able to extract a phylogenetically generalizable signal.

To assess how much genomic information is required for accurate prediction, we retrained and evaluated models on randomly selected feature subsets of varying size, applied consistently to both training and test data, see Table I. Oxygen use, cell envelope, and the mesophile–thermophile binary classification all retain comparable performance even when 60 − 70% of features are removed, including under distant train–test splits, indicating broadly distributed, redundant signal. The minimum completeness required increases with evolutionary distance, reflecting the need for broader genomic coverage to generalize across deep splits. Fine-grained quantitative prediction of OGT requires higher completeness, suggesting it integrates many weak signals rather than a few strongly discriminative markers.”

**Table I:** Genome completeness test. Phenotype predictors were retrained using progressively increasing fractions of randomly selected genes from each genome. The values in the table report the mean ± standard deviation of the minimum fraction of the genome required to reach 95% of the predictive performance achieved using complete genomes. Rows correspond to different phenotypes and evaluation metrics, while columns correspond to different taxonomic holdout levels used for testing. Results are not reported for cases where the predictive performance on complete genomes is low (MCC < 0.5, or *R*^2^ < 0.5).

|  | Random | Family | Order | Class | Phylum | Domain (a-b) | Domain (b-a) |
| --- | --- | --- | --- | --- | --- | --- | --- |
| Oxygen use, MCC | 0.06 ± 0.02 | 0.06 ± 0.02 | 0.07 ± 0.03 | 0.16 ± 0.12 | 0.25 ± 0.18 | 0.62 ± 0.3 | 0.2 ± 0.15 |
| Cell envelope, MCC | 0.07 ± 0.03 | 0.09 ± 0.03 | 0.1 ± 0.05 | 0.32 ± 0.13 | 0.33 ± 0.3 | – | – |
| Sporulation, MCC | 0.06 ± 0.02 | 0.06 ± 0.02 | 0.11 ± 0.14 | 0.65 ± 0.27 | 0.47 ± 0.38 | – | – |
| OGT, MCC | 0.16 ± 0.07 | 0.12 ± 0.07 | 0.33 ± 0.14 | 0.09 ± 0.05 | 0.09 ± 0.06 | – | 0.4 ± 0.35 |
| OGT, $R^2$ | 0.21 ± 0.07 | 0.28 ± 0.11 | 0.53 ± 0.17 | 0.17 ± 0.22 | 0.1 ± 0.05 | – | 0.31 ± 0.22 |
| GC content, MCC | 0.05 ± 0.01 | 0.2 ± 0.19 | 0.05 ± 0.01 | – | – | – | – |
| GC content, $R^2$ | 0.06 ± 0.02 | 0.25 ± 0.13 | 0.06 ± 0.01 | – | – | – | – |

### B Determining the conserved genomic basis of phenotypes

While the genome completeness experiments discussed in Section II-A reveal how much genomic information on average is required for accurate phenotype prediction across evolutionary distances, they do not explain which genes carry the predictive signal or how this signal is distributed across the genome. For annotated data and a trained predictive model, feature selection methods fall into two broad categories: model-specific, and model-agnostic.

A commonly used model-specific approach is SHAP (SHapley Additive exPlanations), see Fig. 4(a), which quantifies a feature’s contribution to prediction by measuring its effect across all possible feature subsets [46]. SHAP identifies the features most strongly used by the model, but reflects model behavior rather than direct biological or causal relationships [47], and can be difficult to interpret when features are correlated [48].

**Figure 4:**
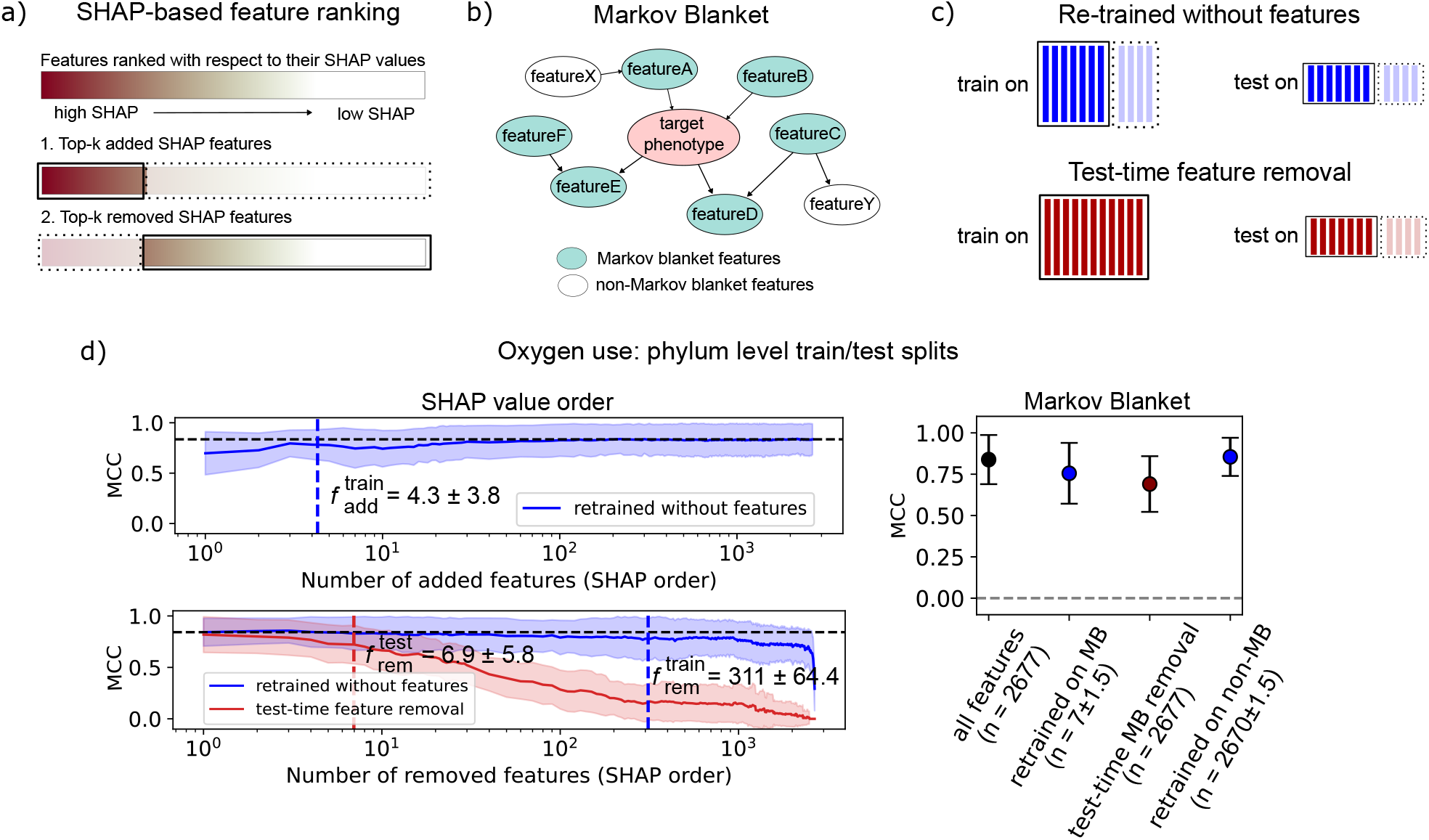
(a) SHAP feature ranking experiments. In the first setting, we train and evaluate a model using the top-k features with the highest SHAP values. In the second setting, we remove the top-k SHAP features and train and evaluate the model on the remaining features.(b) Schematic representation of the Markov blanket in a Bayesian network. The Markov blanket features (blue) informationally shield the target node (pink) from all remaining features (white), such that conditioning on the Markov blanket renders the target independent of the rest of the network. (c) Difference between re-training and test-time feature removal. In re-training, selected features are removed from both the training and test sets, whereas in test-time feature removal, the features are removed only at test time. (d) Oxygen use predictor performance under phylum-level test holdout using selected features. Left: Top-SHAP feature addition and removal experiments. For feature removal, we evaluate two settings: removal during both training and testing, and removal at test time only. Right: Model accuracies obtained using Markov blanket and non-Markov blanket features, compared with the full feature set.

To address these limitations, we additionally apply a model-agnostic method based on Markov blanket (MB) discovery [49]. In this framework, genomic features and the target phenotype are treated as nodes in a Bayesian network, see Fig. 4(b), and the MB is defined as the minimal set of features that renders the target conditionally independent of all others:

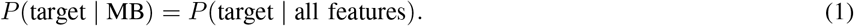

Together, these methods allow us to move beyond model accuracy toward biological insight. Features identified by both SHAP and the Markov blanket represent high-confidence, conditionally indispensable signals, whereas SHAP-only features likely reect predictive but redundant correlations, and Markov-blanket-only features capture weak yet essential dependencies underexploited by the model.

To assess the robustness and minimality of the feature sets identified by SHAP, we perform a controlled feature ranking and reduction analysis. For each train/test data split, we rank all features according to their SHAP values and sort them from the most SHAP-important to the least important, see Fig. 4(a). Then we re-train and re-evaluate the model on the reduced feature set in two ways, see Fig. 4(c), top. In the first strategy, features are added sequentially in order of decreasing SHAP importance. This procedure identifies the minimal set of top SHAP-ranked features required for the predictor to reach 95% of the MCC obtained with the full training feature set. In the second strategy, features are removed sequentially, starting from the most important ones, which identifies the maximal set of top SHAP-ranked features that can be removed during training while maintaining at least 95% of the MCC obtained with the full training feature set. We denote these quantities by 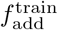 and 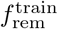 for feature addition and feature removal, respectively. In addition, we perform a test-time feature removal experiment, see Fig. 4(c), bottom. In this setting, the model is trained on the full feature set but evaluated on progressively reduced feature sets obtained by removing features in order of decreasing SHAP importance. Feature removal is applied at test time only by erasing their information, implemented by setting the corresponding count values to NaN in XGBoost [50] (see Supplementary Materials for details). Analogously, we determine the maximal number of top SHAP-ranked features that can be removed at test time while maintaining at least 95% of the MCC obtained with the full training feature set, and denote this quantity by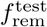

Finally, we identify the MBs for each phenotype and each data fold, and evaluate model performance using the MB feature sets in a similar manner to the SHAP-based analysis, see Fig. 4(d), right. Corresponding plots for the remaining phenotypes, analogous to Fig. 4(d,e), are provided in the Supplementary Materials. The Markov blanket sizes and the SHAP-derived statistics 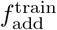 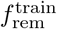 and 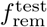 across all phenotypes are summarized in Table II.

**Table II:** Markov blanket (MB) sizes and SHAP-based feature counts across phenotypes. 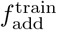 denotes the minimum number of top SHAP-ranked features that must be added during training to reach 95% of the MCC obtained with the full training feature set. 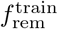 denotes the maximum number of top SHAP-ranked features that can be removed during training while maintaining at least 95% of the MCC achieved with the full training feature set. 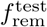 denotes the maximum number of top SHAP-ranked features that can be removed only at test time while maintaining at least 95% of the MCC achieved with the full training feature set. Values represent the mean *±* standard deviation across phylum-level train–test splits.

|  | Oxygen use | Cell envelope | Sporulation | OGT (binary) | GC content (binary) |
| --- | --- | --- | --- | --- | --- |
| $f_{\text{add}}^{\text{train}}$ | $4.3 \pm 3.8$ | $2.3 \pm 1.9$ | $13.4 \pm 7.1$ | $98.1 \pm 13.2$ | $44.7 \pm 17.2$ |
| $f_{\text{rem}}^{\text{train}}$ | $311 \pm 64.4$ | $1095 \pm 71$ | $1159.3 \pm 141.3$ | $1539.3 \pm 114$ | $729 \pm 71.1$ |
| $f_{\text{rem}}^{\text{test}}$ | $6.9 \pm 5.8$ | $2 \pm 0.1$ | $3.2 \pm 0.9$ | $114.3 \pm 28$ | $79.5 \pm 21.7$ |
| MB size | $7 \pm 1.5$ | $4.3 \pm 0.9$ | $7.4 \pm 2.5$ | $22.3 \pm 2.8$ | $19.1 \pm 2.5$ |

For oxygen use, both 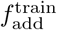 and the MB size are small, indicating that most predictive performance can be recovered from only a few features. However, prediction accuracy remains high after removing hundreds of top-SHAP features or retraining on non-MB features alone, revealing substantial redundancy in the genomic signal. Despite this, the predictor relies strongly on a small subset of informative features, as their removal at test time causes a rapid decline in performance. Cell envelope structure and sporulation show a similar pattern of distributed signal coupled with strong dependence on a few highly informative features. In contrast, OGT and GC content rely on a broader feature set, allowing more features to be removed before performance deteriorates.

Finally, oxygen use, cell envelope, and sporulation have 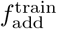 values smaller than their MB sizes, indicating that a small subset of features is sufficient for accurate prediction, while the full Markov Blanket captures a broader set of statistical dependencies with the target variable. Conversely, the smaller MB sizes of OGT and GC content relative to 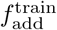 suggest that many top SHAP-ranked features are redundant.

We next examined the composition of the MB and top-SHAP feature sets for each phenotype. To obtain robust feature lists, we retained the features most frequently identified across phylum-level train–test folds. We then quantified the association between COG presence and phenotype occurrence within each phylum using a signed enrichment score (Eq. 6); positive values indicate enrichment in phenotype-positive organisms, and negative values indicate enrichment in phenotype-negative organisms. For oxygen use, the most important features and their correlations with phenotype occurrence are shown in Fig. 5. We find that ferredoxin oxidoreductase (COG1014), rubrerythrin (COG1592), and anaerobic ribonucleoside-triphosphate reductase (COG1328) are strong predictors of anaerobic metabolism, whereas heme A synthase (COG1612) and heme O synthase (COG0109) are strong predictors of aerobic metabolism. These results are consistent with previous observations [14] and are readily interpretable in terms of known biology: the strongest predictors of anaerobic metabolism are hallmark enzymes of anaerobes, while the strongest predictors of aerobic metabolism are involved in biogenesis of the aerobic electron transport chain. In addition, phosphoribosylaminoimidazole carboxylase (COG0026) and type II dihydrodipicolinate synthase (COG1002) appear as predictors of aerobic metabolism, and a two-component system response regulator (COG1350) also emerges as an aerobic predictor that, to our knowledge, has not been highlighted previously in this context. Analogous plots for other phenotypes can be found in Supplementary Materials, Fig. 10 for sporulation, Fig. 11 for optimal growth temperature, and Fig. 14 for GC content.

**Figure 5:**
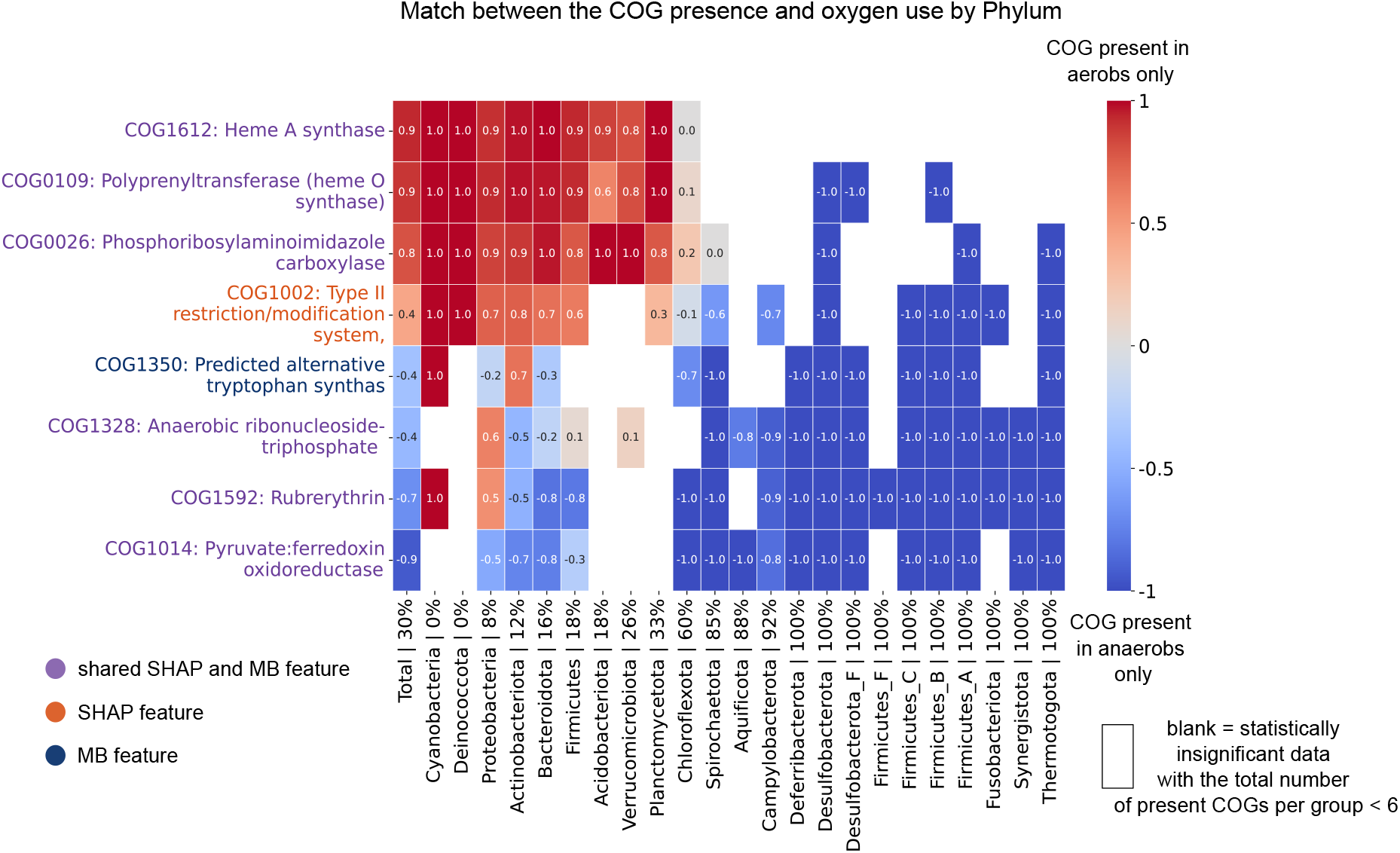
Top SHAP-ranked features and Markov blanket (MB) features associated with the oxygen use phenotype, and their presence–phenotype pattern overlap across phyla. Feature colors indicate membership in the top-SHAP set (orange), the MB (dark blue), or their intersection (purple). The score quantifies the overlap between COG presence and phenotype occurrence (see Methods). A score of +1 (red) indicates perfect agreement between COG presence and oxygen use, whereas a score of −1 (blue) indicates perfect agreement between COG presence and and the anaerobic (phenotype-negative) class, i.e. the COG is present exclusively in anaerobic organisms.

The most important gene families for the cell envelope predictor also make biological sense. The 8 genes in the top-SHAP and Markov blanket set include core components of outer membrane biogenesis and maintenance, including genes for lipopolysaccharide synthesis and export to the outer membrane (COG0763, LpxB - lipid-A-disaccharide biosynthesis; COG0795, LptF/G - LPS export; COG0767, MlaE - intermembrane phospholipid transport), genes involved in cell wall biosynthesis (COG5009, membrane-bound penicillin-binding protein) along with diderm-specific genes that localise to the periplasm or outer membrane such as COG1706, FlgI - flagellar basal body P-ring protein, see Supplementary Fig. 9. Interestingly, the most important genes are either specific to diderms or mainly found in diderms. This finding suggests that didermy is specified by a defined, common genetic toolkit that is conserved across diderm lineages, consistent with previous work [51], [35]. By contrast, monoderms appear to be characterised by an absence of the diderm toolkit rather than the presence of monoderm-specific genes. Given the interleaving of monoderm and diderm lineages on the bacterial phylogeny, these results are consistent with the hypothesis that monodermy evolved convergently, and on different genetic backgrounds, on multiple occasions [51].

As in the case of the phenotypes discussed above, the SHAP and Markov Blanket analyses identified many expected gene families with a known biological link to the trait of interest. However, each of these traits also had a larger number of relevant features than in the case of oxygen use or cell envelope architecture, and - while performance was competitive with existing approaches - were generally less well predicted by the best-fitting models. Unsurprisingly, the most important gene families for predicting sporulation were those directly involved in the process, including spore coat (COG5577), maturation (COG2715) and control (COG4326) proteins, among others, see Supplementary Fig. 10. Top-ranked gene families for predicting thermophily included the hallmark enzyme of hyperthermophiles, reverse gyrase (COG1110) (add refs to reverse gyrase papers and its use as hallmark), an enzyme involved in DNA repair of 8-oxoguanine mutations (COG4047) that are elevated at high temperatures [52] along with other proteins including tRNA modification enzymes (COG1590, COG1818) that have previously been implicated in thermoadaptation [53], see Supplementary Fig. 11. The top-ranked genes for predicting high-GC organisms were somewhat more difficult to interpret. Several of the top-ranked genes are involved in DNA repair (COG4544), including in the context of UV damage (COG1533), or trehalose biosynthesis (COG1877, COG3280); these are all functions that are important for high-GC soil bacteria (e.g. Actinomycetota), which experience a range of environmental stressors. In this case, it may be that the predictive power comes from the indirect association between high-GC genomes and soil ecology, rather than a direct link between the predictive gene families and the trait.

### C. Ancestral phenotype prediction

A key motivation for representing genomes as gene family copy numbers is that ancestral copy number profiles can be inferred using phylogenetic reconciliation methods [26], enabling phenotype prediction beyond extant taxa [14]. However, ancestral genomes are reconstructed probabilistically rather than observed directly, and inferred gene content becomes increasingly uncertain toward the root due to cumulative gene duplications, transfers, and losses [54]. Robust phenotype predictors must therefore tolerate substantial noise and incompleteness in genomic features.

Training on noise-augmented data improves robustness by exposing models to controlled perturbations, encouraging noise-invariant representations that are less sensitive to feature corruption at inference time [55], [56], [57], [58]. We augmented the training data with synthetic noisy genomes. For each genome, multiple noisy replicas were generated by introducing false-negative and false-positive gene counts, simulating gene loss and spurious gene presence that can arise during genome assembly and annotation, see Fig. 6 (a). The noise levels were sampled from predefined distributions to capture a range of possible error regimes. For each phenotype, we evaluated multiple augmentation configurations and selected the scheme that maximized predictive performance on noisy test datasets. Full details of the noise model and parameter selection procedure are provided in Section 8 in Supplementary Materials.

**Figure 6:**
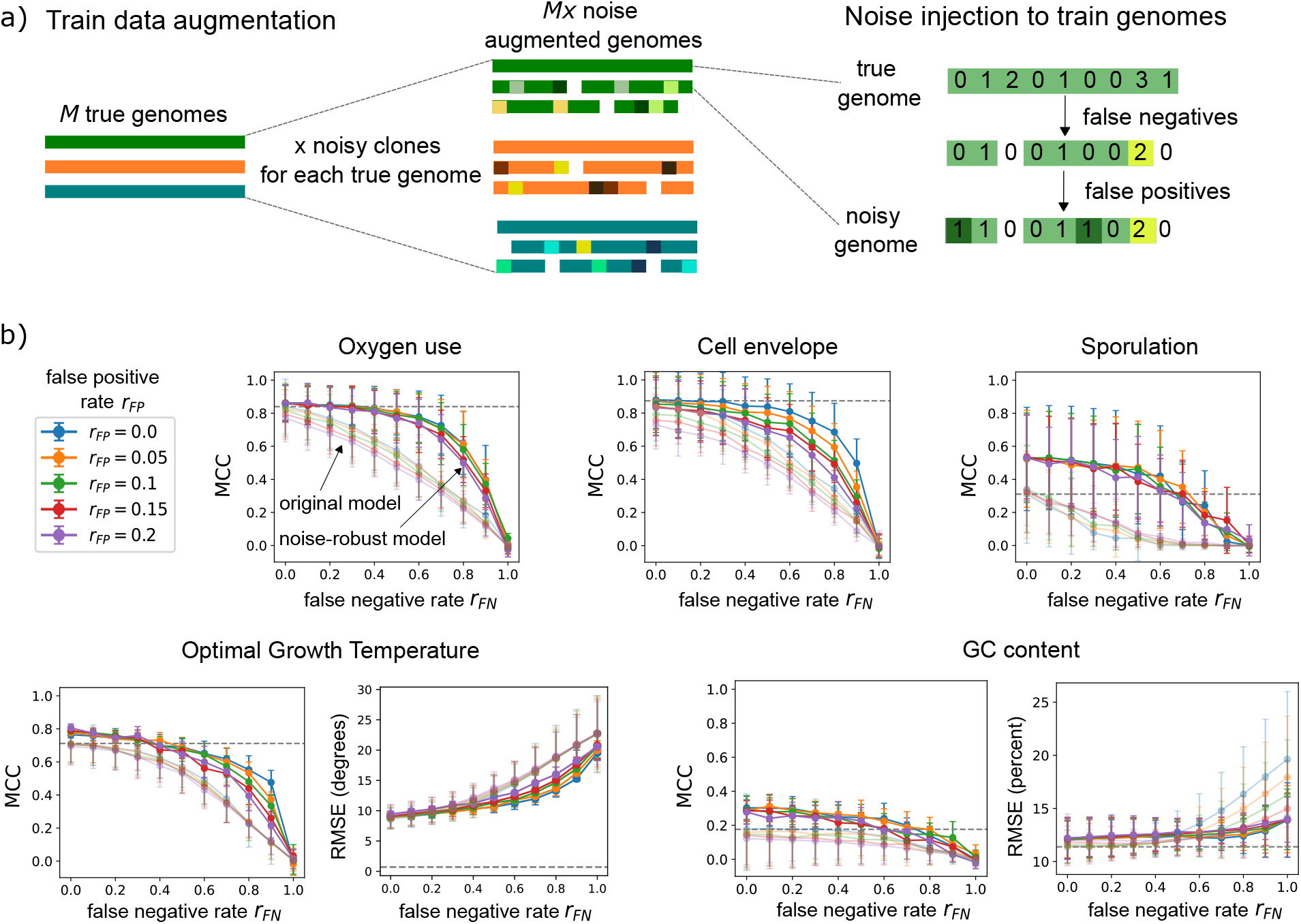
a) Training data augmentation scheme. Each genome is augmented with *x* noisy replicates, where noise is introduced by injecting false-positive and false-negative COG counts. b) Performance comparison between the original models trained on noise-free datasets (semi-transparent lines) and the best noise-robust models trained on noise-augmented datasets (solid lines). Prediction accuracy is shown across different false-positive (*r*_*FP*_) and false-negative (*r*_*FN*_) noise rates applied to the test data.

For oxygen use and cell envelope phenotypes, we observe a clear improvement in model accuracy across the full range of noise levels in the test datasets, see Fig. 6 (a,b). For the sporulation phenotype, this approach not only improves performance under high noise conditions but also acts as an effective regularization mechanism, leading to improved accuracy even on nearly noise-free test data, see Fig. 6(c). For optimal growth temperature, we observe a modest performance improvement without compromising baseline performance at low noise levels, see Fig. 6(d). In contrast, for GC content prediction, gains at higher noise levels are accompanied by a slight reduction in performance at low noise levels, indicating a trade-off between robustness and accuracy in low-noise regimes, see Fig. 6(e).

Next, we applied the best-performing models to ancestral phenotype inference. Our predictions suggest that the last bacterial common ancestor (LBCA) was likely an anaerobic, non-sporulating, moderately thermophilic diderm, see Fig. 7(a-e) and Supplementary Table 7. The models differ in prediction confidence across the ancestral timescale, with cell envelope showing the highest confidence and GC content the lowest, see Fig. 7(f). For GC content, ancestral states therefore cannot be inferred with comparable confidence. This is consistent with its lineage-restricted genomic signal identified in Section II-B: even after noise augmentation, the GC content predictor recovers only modest accuracy on noisy genomes, see Fig. 6(e) and already degrades sharply at class- and phylum-level holdouts in extant data, see Fig. 3; Table I, suggesting that noise augmentation can reinforce a redundant signal but cannot recover one whose informative components are inherently lineage-specific.

**Figure 7:**
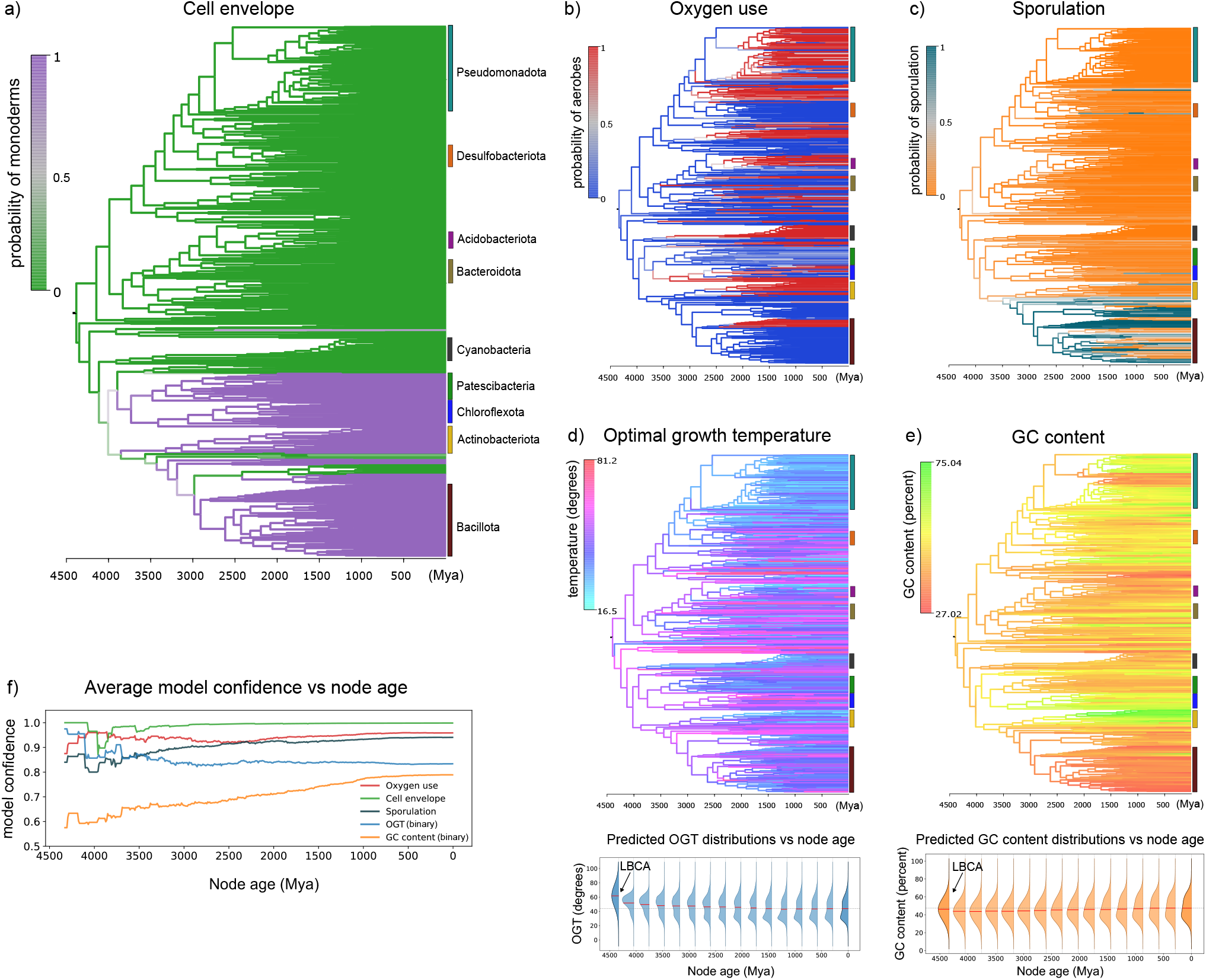
Ancestral phenotype prediction for the bacterial species tree from [14]. Branch colors represent the predicted class probability for the binary classifiers (a-c) and the final predicted value for the continuous regressors (d-e). (a) Cell envelope.(b) Oxygen use. (c) Sporulation. (d) Top: Optimal growth temperature tree. Bottom: predicted OGT distributions across nodes binned by age; the red line indicates the mean of each distribution and the dashed horizontal line corresponds to the predicted mean OGT of extant genomes in the dataset. (e) Top: GC content tree. Bottom: predicted GC content distributions across nodes binned by age, as in (d). (f) Average prediction confidence of the binary classifiers as a function of node age for the phenotypes shown in (a-c).

## III. Discussion

Overall, our results suggest that many bacterial phenotypes of evolutionary interest can be inferred from gene content reconstructions back to the root of the bacterial domain, because information pooled from multiple phenotype-relevant genes can help to overcome the uncertainty inherent in single gene reconstructions. To understand why some ancient phenotypes can be predicted and some cannot, we investigated how their genomic signal is structured and how well prediction generalizes across evolutionary distances. We found that oxygen use and cell envelope structure are underpinned by broadly conserved, genome-wide signals: even after removing the majority of genomic features, predictive performance is largely retained, and models generalize well even under phylum- and domain-level holdout splits. Sporulation, OGT, and GC content, by contrast, show sharper performance declines with evolutionary distance, consistent with more lineage-restricted or environmentally contingent genomic encoding. At least part of the explanation may be that cell envelope architecture and oxygen metabolism are directly implemented by specific gene families (for example, those encoding components of the aerobic electron transport chain, or the enzymes that produce outer-membrane lipopolysaccharides and their transporters). By contrast, organisms with particular OGTs or genomic GC contents may tend to possess certain gene families (enabling prediction over shorter evolutionary distances), but these properties are not, as far as is currently known, directly “caused” by a specific set of genes. This explanation does not account for the relatively narrow predictive power of the sporulation classifier, since this trait is also encoded by specific gene families. However, unlike cell envelope and metabolic phenotypes, sporulation is found in only a handful of modern bacterial lineages, which may explain its different behaviour in our analyses.

Despite the broad signal availability for at least some of these traits, our test-time feature removal experiments reveal an important asymmetry: models tend to exploit a small subset of highly informative features rather than the full distributed signal. When these features are missing or corrupted at inference time (as they inevitably are in reconstructed ancestral genomes), performance drops sharply even though alternative informative features exist. This distinction between signal availability and signal usage has practical consequences for ancestral inference and motivates our noise augmentation strategy. By exposing models to controlled false-positive and false-negative perturbations during training, we encourage reliance on redundant, distributed signals rather than fragile feature subsets, substantially improving robustness across the full range of noise levels. Crucially, however, noise augmentation can only reinforce signal that is already broadly distributed: for GC content, whose predictive signal is inherently lineage-specific, augmentation improves robustness at high noise but ancient GC content cannot be inferred.

After noise augmentation, the models stay well calibrated even in the presence of substantial false-positive and false-negative noise. Their predicted class probabilities remain meaningful on the kind of incomplete, error-prone feature vectors produced by ancestral genome reconstruction. Because expected gene copy numbers can be inferred at internal nodes by phylogenetic reconciliation [26], this combination lets us carry prediction — and a calibrated estimate of its confidence — back along the bacterial tree to its root. We exploited this by tracking model confidence as a function of node age, from extant tips to the last bacterial common ancestor (LBCA), see Fig. 7(f). For phenotypes underpinned by broadly conserved, genome-wide signal, confidence decays only slowly toward the root and remains high at the deepest nodes, whereas for lineage-restricted phenotypes such as GC content it falls to near-random well before the LBCA. The practical consequence is that some phenotypes can be reconstructed confidently into the Archaean.

Applying our pipeline to a species tree of 1007 bacteria, we infer that the LBCA was likely an anaerobic, non-sporulating, moderately thermophilic diderm, with the cell envelope and oxygen-use predictions in particular retaining high confidence at the root, consistent with and extending recent work linking genomic signatures to early Earth history [14]. These ancestral predictions are broadly consistent with current hypotheses about early bacterial evolution. The inference of an anaerobic LBCA agrees with the prevailing view that most bacterial lineages predate the Great Oxidation Event and with the conclusions of [14].

The inference that the LBCA and the ancestors of Gracilicutes and Terrabacteria were diderms agrees with a growing body of work suggesting that the double membrane is an ancestral bacterial feature and that monodermy represents a derived state [51], [59], [60], [61]. The genes most predictive of didermy in our analysis overlap with the canonical diderm machinery — lipid A and lipopolysaccharide (LPS) biosynthesis and outer-membrane assembly (Section II-B) — consistent with previous candidate-gene analyses [62], [63], supporting the biological validity of the learned signal. The classifier further predicts multiple independent losses of the outer-membrane machinery within Terrabacteria, including on the lineages leading to Patescibacteria and to Firmicutes, in agreement with recent reconstructions [51], [60], [61]. The best-supported scenario is therefore one in which the double membrane is ancestral in Bacteria and was lost convergently on multiple independent lineages — a conclusion that stands in contrast to earlier views of early bacteria as monoderm organisms resembling modern Firmicutes or Actinobacteria [64], [65], and that highlights the outer membrane as an under-appreciated distinguishing feature between early Bacteria and Archaea.

Although the precise optimal growth temperature of the LBCA cannot be inferred with high confidence, our analysis is broadly consistent with previous work suggesting a thermophilic LBCA [66], [67], with optimal temperatures in the 46–75°C range (Fig. 7). In more recent times, Bacteria adapted to somewhat cooler conditions, with some signal of bimodality and a peak emerging at 37°C. This might result from adaptation to eukaryote-associated lifestyles - including most recently human guts - although it may also reflect the nature of how experimental OGT data is collected, with 37°C a frequently-tested value [68].

While our results encourage optimism about the detail with which ancient life can be reconstructed given modern data, several limitations should be noted. Ancestral gene content is reconstructed probabilistically, and expected copy numbers at deep nodes carry substantial uncertainty that compounds toward the root. Our noise augmentation scheme simulates this uncertainty but cannot fully capture its phylogenetic covariance structure. Published experimental information on phenotypes collated in BacDive and related databases is highly valuable but incomplete and unevenly distributed across lineages, which may bias both training and the interpretation of ancestral predictions. In addition, our pipeline was evaluated on five phenotypes chosen to span a range of biological and predictive complexity; how well these findings generalize to other traits, particularly those with more complex polygenic architectures or stronger environmental contingency, remains to be explored.

Beyond ancestral inference, the framework developed here offers a general approach for probing the evolutionary signal of any phenotype encoded in genomic gene content, and for identifying new gene families that appear to be associated with phenotypes of interest. The combination of taxonomy-aware evaluation, feature redundancy analysis, and noise-robust training provides a principled basis for assessing which phenotypes are tractable targets for deep-time reconstruction and which are not— a question of broad relevance as ancestral genome reconstruction methods continue to improve. An immediate extension would be to apply the same pipeline to other phenotypes of evolutionary and ecological interest, such as nitrogen fixation, photosynthesis, or antibiotic resistance, and to use the resulting ancestral predictions as calibration points in molecular clock analyses, as demonstrated previously [14].

## IV. Methods

### A. Two stage OGT and GC content prediction

OGT values are heavily concentrated between 20 and 40°C, see Fig. 3(a), left. To predict continuous OGT, we employ a two-stage gating approach. An XGBoost classifier first partitions genomes into mesophiles (OGT ≤ 45°C) and thermophiles (OGT ≥ 45°C), yielding probabilities *p*_meso_ and *p*_thermo_. Two separate XGBoost regressors are then trained on each group, producing predictions OGT_meso_ and OGT_thermo_. The final OGT estimate is their probability-weighted combination:

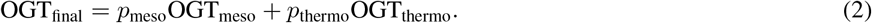

The GC annotation distribution in our dataset is bimodal, with two prominent peaks around 38% and 62%, suggesting the presence of two major genomic composition groups, see Fig. 3(b), left. Similarly to the OGT, we perform the binary classification, and split the data into the low GC content (≤ 50), and high GC content (≥ 50) categories:

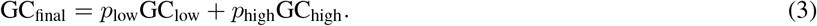

### B. Phylogeny-based phenotype prediction

In this section, we describe the algorithm used for phenotype prediction based solely on phylogenetic relatedness. We reused the same 30 train–test splits generated for the XGBoost predictor at each taxonomic level. In each split, 80% of the genomes constitute the training set and are treated as annotated reference genoms, while the remaining 20% form the test set whose phenotypes must be inferred.

For each test genome, we traverse the phylogenetic tree to identify the nearest ancestral clade whose descendant leaves belong entirely to the training set. The taxonomy-aware splits ensure that such a clade exists for every test genome. The phenotype of the test genome is then inferred from the phenotypes of the genomes within this closest training clade. For continuous phenotypes, the prediction is defined as the mean phenotype value across the clade. For binary phenotypes, the predicted class corresponds to the majority class among the genomes in the clade. We note that at the domain level, phenotypes of bacterial genomes are inferred as the average (or majority class) of archaeal phenotypes, and vice versa.

For random train–test splits, where such taxonomic separation is not guaranteed, we instead assign the phenotype of the single closest training genome identified using the pairwise phylogenetic distance matrix derived from the genome tree of the GTDB, see Fig. 8.

**Figure 8:**
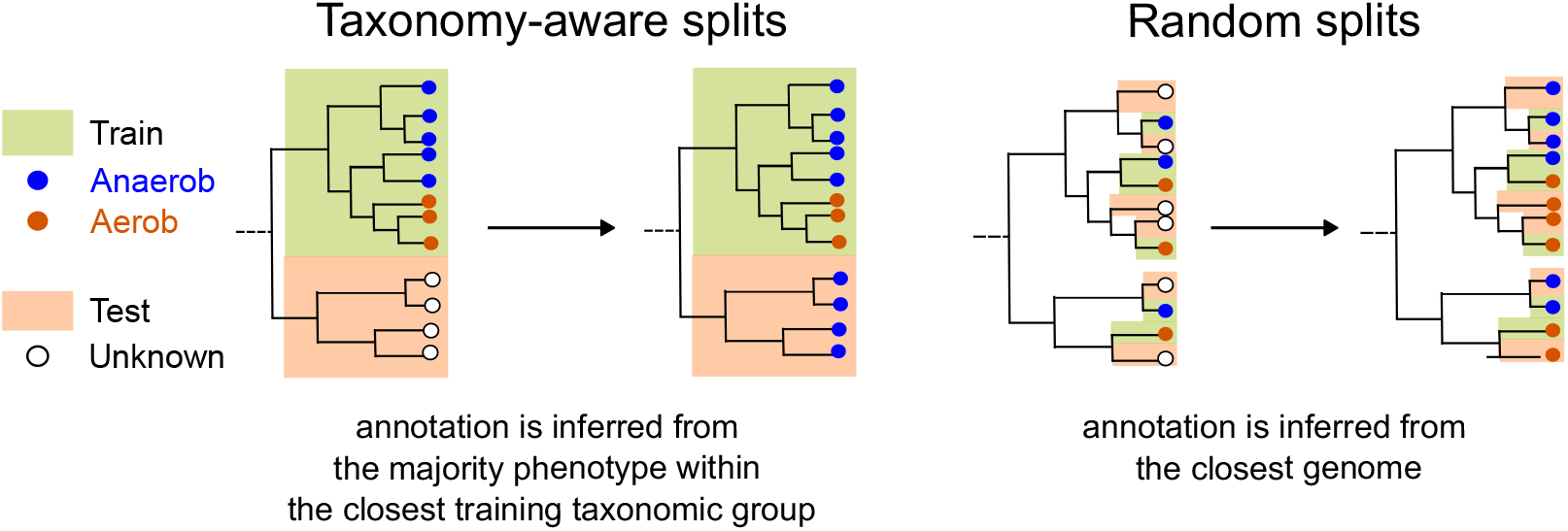
Phylogeny-based annotation predictions. The training data (80% of genomes) are treated as known annotations from which the remaining 20% are inferred. In taxonomy-aware train/test splits, the phenotype of each test genome is inferred as the majority phenotype in the binary case or the mean phenotype in the continuous case among genomes belonging to the closest taxonomic group contained in the training set. In random train/test splits, the phenotype is inferred from the single closest genome.

### C. Feature selection: Pattern overlap score

Let *y*_*i*_ ∈ {0,1} encode the phenotype of genome *i* (e.g. 0 = anaerobe, 1 = aerobe; 0 = diderm, 1 = monoderm etc). Let *g*_*i*_ ∈ {0,1} denote the presence or absence of a given COG in the genome *i*. We define the pattern overlap as

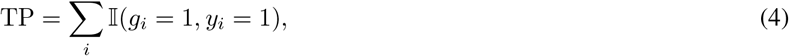

which counts the number of genomes in which both the COG and the phenotype encoded by 1 are present. Conversely, we define the pattern mismatch as

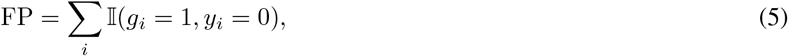

which counts the number of genomes that contain the COG but exhibit the phenotype encoded by 0.

Combining Eq. 4, and Eq. 5, we define the final pattern overlap score

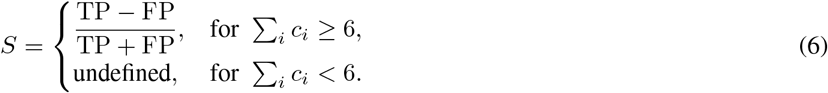

We note that *S* is not defined for COGs with fewer than six total occurrences across all genomes, as such cases lack sufficient statistical support.

### D. Noise-based training data augmentation

We augment the train datasets by generating *x* noisy clones of each true genome 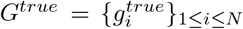 where 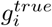 denotes the true count of gene *i*. Noise is introduced by injecting false negative and false positive counts as follows.

- To model false negatives, for each gene count *g*_*i*_, we subtract a Poisson-distributed number of counts proportional to 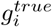

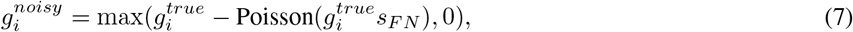

where *s*_*FN*_ denotes the false negative rate.
- False positives are then introduced for genes with zero observed counts by adding Poisson-distributed counts:

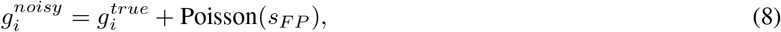

where *s*_*FP*_ is the false positive rate.

Together, this procedure simulates gene loss and spurious gene presence, approximating the uncertainty induced by genome reconstruction errors while preserving the overall structure of the genome.

Rather than fixing the false negative *s*_*FN*_ and false positive *s*_*FP*_ rates, we consider stochastic noise regimes in which these rates are independently sampled from predefined probability distributions for each noisy genome. Specifically, we focus on three distribution families: uniform, exponential, and gamma. Each distribution is parameterized by its mean value, denoted *λ*_*FN*_, and *λ*_*FP*_ for false negatives and false positives, respectively. Each noise augmentation scheme is thus defined by the choice of the three parameters (*x, λ*_*FN*_, *λ*_*FP*_). For each phenotype, we systematically explore combinations of these parameters, retrain XGBoost models for each configuration, and evaluate performance gains on noisy test datasets with varying noise levels. The test-time noise is introduced by setting non-zero gene counts to zero at rate *r*_*FN*_ and incrementing all counts by one at rate *r*_*FP*_. We then quantify model robustness across the full test-noise range by computing the area under the MCC-versus-noise curves, similar to those shown in Fig. 6. Finally, for each phenotype, we select the augmentation scheme that maximizes the average MCC curve integral, corresponding to the greatest overall robustness improvement on noisy test data. The performance improvements of the best-performing models for each phenotype are summarized in Fig. 6 (see Section 8 in Supplementary Materials for additional details on model selection and augmentation configurations).

## V. Data and code availability

All code is available at https://github.com/oist/gene-context. Data and trained models are deposited at Zenodo (DOI: 10.5281/zenodo.21018005).

## VI. Acknowledgments

We are grateful for the help and support provided by the Scientific Computing and Data Analysis section of Core Facilities at OIST. Our research is funded by the John Templeton Foundation (63451 to T.A.W., G.Sz. and A.S.); the opinions expressed in this publication are those of the authors and do not necessarily reflect the views of the John Templeton Foundation.

