## Supplementary Materials for "Models trained with noisy genomes extend bacterial phenotype prediction into deep time"

### 1 tSNE embeddings of the datasets

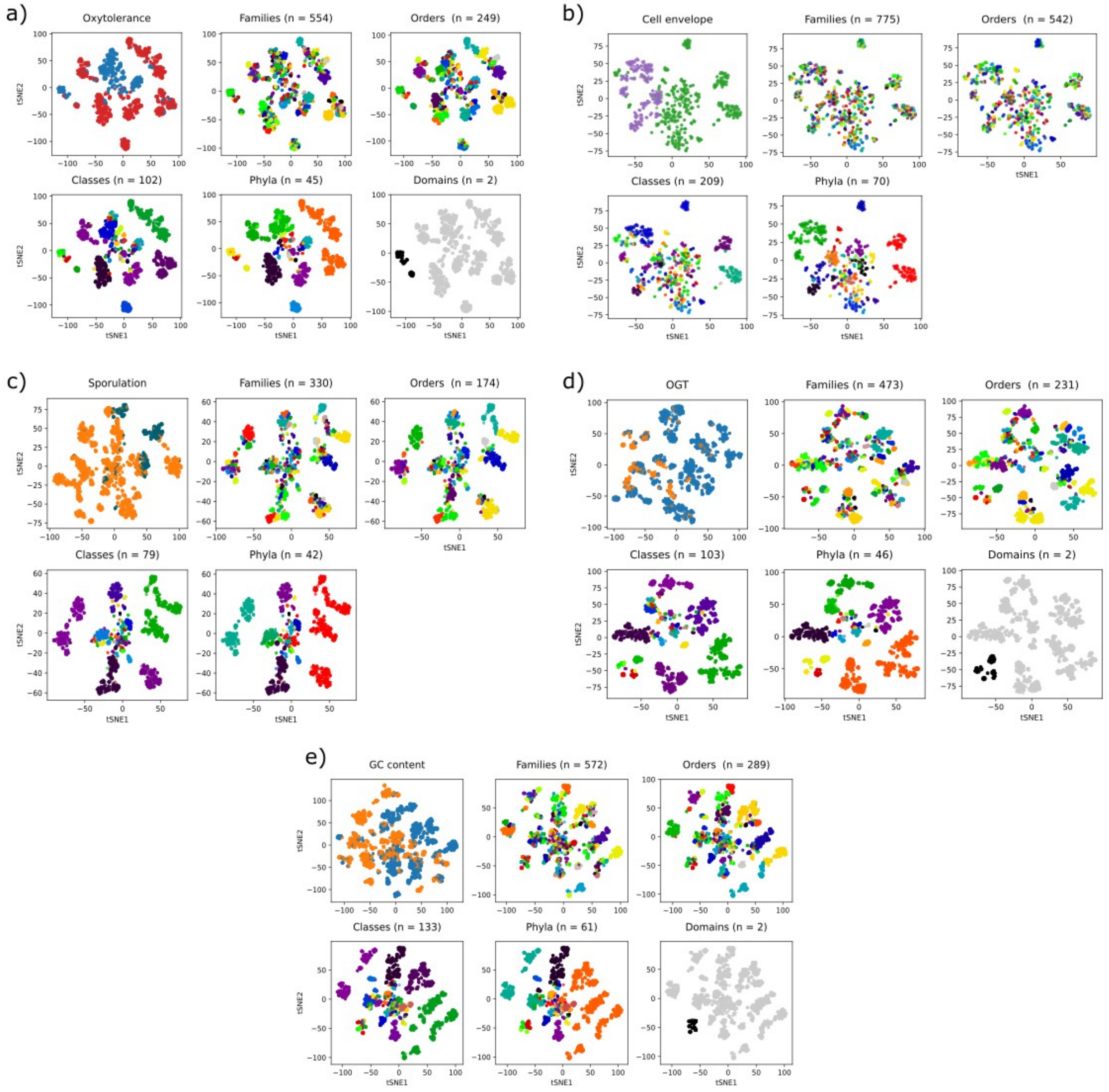


Figure 1: tSNE embeddings for the phenotype datasets colored with respect to the phenotype annotations and taxononomic groups. (a) Oxytolerance. (b) Cell envelope. (c) Sporulation. (d) Optimal growth temperature (OGT). (e) GC content.

#### 2 Phenotype similarity across taxonomy levels

We explore phenotype similarity within taxonomic groups to understand how phenotypic signal decays with evolutionary distance. For each taxonomic group, we compute the probability that a randomly sampled pair of individuals shares the same phenotype. For a binary phenotype, consider a taxonomic group of nn $n$ individuals, of which $n_0$ have phenotype 0 and $n_1$ have phenotype 1, so that $n_0 + n_1 = n$. The probability that a randomly sampled pair is concordant (i.e., both 0 or both 1) is:

$$P_{\text{taxa}} = \frac{\binom{n_0}{2} + \binom{n_1}{2}}{\binom{n}{2}}.$$

To assess whether this concordance exceeds what would be expected by chance, we compute the same quantity for the full dataset:

$$P_{\text{baseline}} = \frac{\binom{N_0}{2} + \binom{N_1}{2}}{\binom{N}{2}},$$

where $N$ is the total number of genomes, and $N_0$ and $N_1$ are the total counts of phenotype 0 and 1 respectively. We then define the phenotypic homogeneity index as:

$$H = \frac{P_{\text{agree}} - P_{\text{baseline}}}{1 - P_{\text{baseline}}}. \tag{1}$$

$H$ measures the excess concordance within a taxonomic group relative to the global baseline, normalized to the $[0, 1]$ range. $H = 0$ indicates that phenotype concordance within the group is no greater than expected from the global phenotype frequencies alone, while $H = 1$ indicates perfect phenotypic homogeneity within the group. Negative values indicate that the group is more phenotypically diverse than the global baseline, i.e. it is enriched for discordant pairs.

For continuous phenotypes, we compute the mean absolute difference in phenotype values between randomly sampled pairs of individuals, averaged over all pairs within each taxonomic group. The phenotypic homogeneity index $H$ (for binary phenotypes) and the mean absolute difference (for continuous phenotypes) are reported across all phenotypes and taxonomic levels in Figs. 2 and 3.

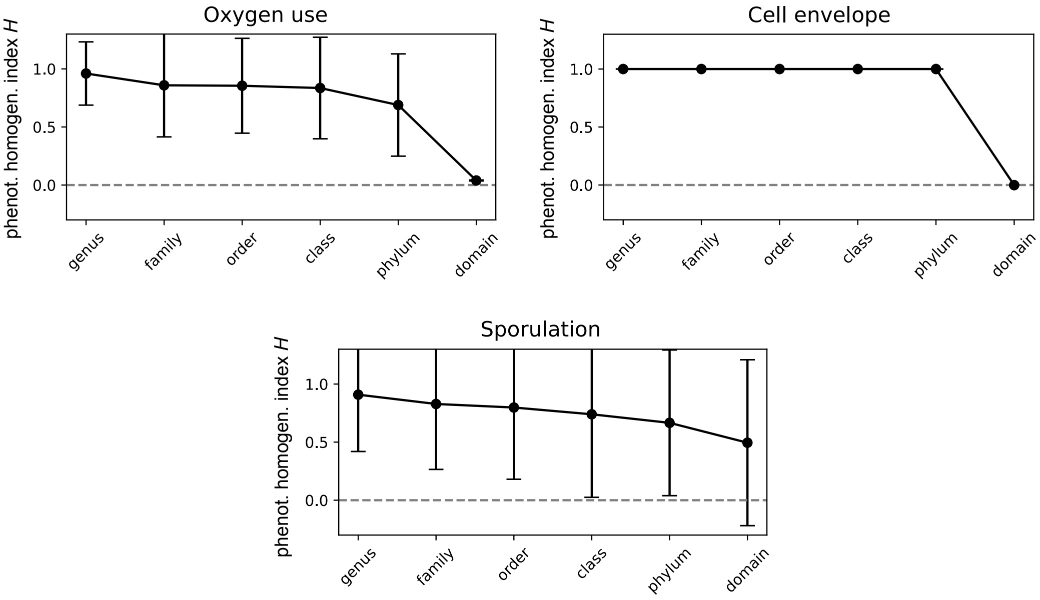


Figure 2: Phenotypic homogeneity index $H$ across taxonomic levels for oxygen use, cell envelope, and sporulation phenotypes.

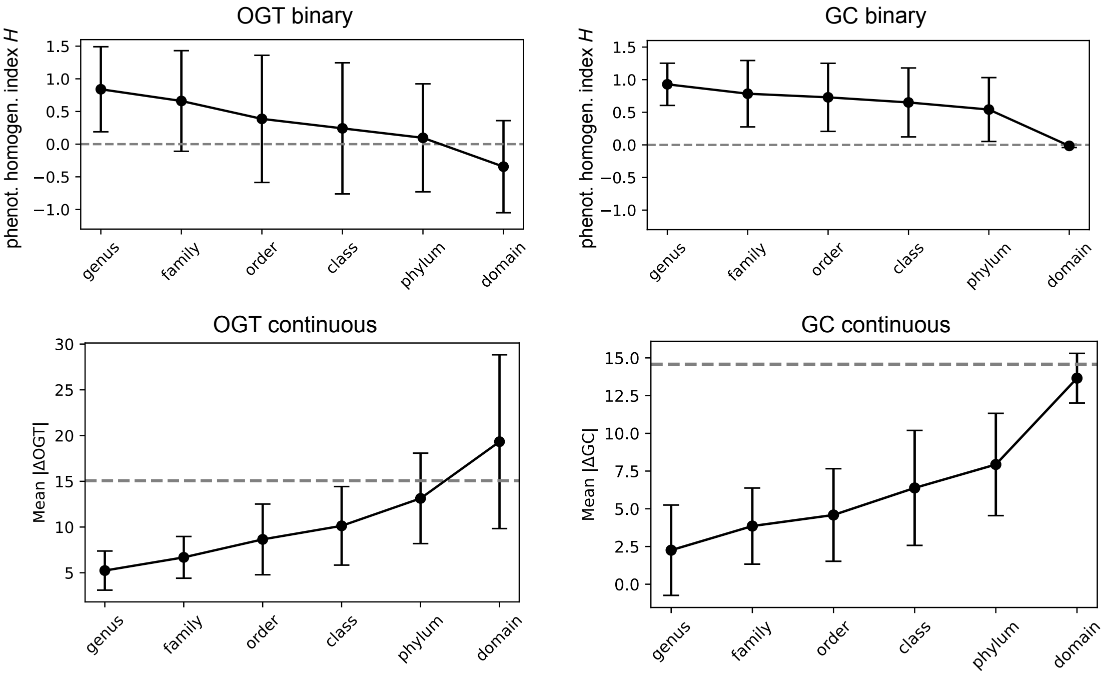


Figure 3: Phenotypic homogeneity index $H$ and the mean absolute difference across taxonomic levels for OGT and GC content phenotypes.

### 3 Model Calibration and Uncertainty Quantification

#### 3.1 Classifier Calibration: Expected Calibration Error

The binary classifier assigns each node a probability $p \in [0, 1]$ of belonging to a class. A well-calibrated classifier should satisfy

$$P(y = 1 \mid \hat{p} = p) = p$$

for all $p$, i.e. among all predictions with confidence $p$, the true positive rate should equal $p$. We quantify deviation from this ideal using the Expected Calibration Error (ECE) [**?**]:

$$\text{ECE} = \sum_{m=1}^{M} \frac{|B_m|}{n} \left|\text{acc}(B_m) - \text{conf}(B_m)\right|, \tag{2}$$

where predictions are grouped into $M$ equally spaced bins $B_m$ by confidence, $|B_m|$ is the number of predictions in bin $m$, $n$ is the total number of predictions, $\text{acc}(B_m)$ is the fraction of correct predictions in the bin, and $\text{conf}(B_m)$ is the mean predicted probability in the bin. ECE ranges from 0 (perfect calibration) to 1 (maximally miscalibrated).

#### 3.2 Regression Calibration: Mean Quantile Calibration Error

The regression component of the gating model produces, for each node, a point prediction $\hat{y}$ alongside an 80% prediction interval $[\hat{Q}_{0.1}, \hat{Q}_{0.9}]$ estimated via quantile regression at the 10th and 90th percentiles. A well-calibrated quantile regressor should satisfy

$$P(y \leq \hat{Q}_\tau) = \tau$$

for each target quantile $\tau$. We quantify deviation from this using the Mean Quantile Calibration Error (MQCE):

$$\text{MQCE} = \frac{1}{|\mathcal{T}|} \sum_{\tau \in \mathcal{T}} \left| \frac{1}{n} \sum_{i=1}^{n} \mathbf{1}[y_i \leq \hat{Q}_\tau(x_i)] - \tau \right| \tag{3}$$

where $\mathcal{T} = \{0.1, 0.9\}$ is the set of target quantiles, $\hat{Q}_\tau(x_i)$ is the predicted $\tau$-quantile for observation $i$, and $\mathbf{1}[\cdot]$ is the indicator function. MQCE ranges from 0 (perfect calibration) to 1 (maximally miscalibrated), and is the natural regression analog of ECE.

#### 3.3 Prediction Uncertainty: Combined Variance for gating models

For each test genome, we quantify total prediction uncertainty via the law of total variance applied to the full gating pipeline. Let $p$ denote the classifier's probability of assigning a node to class 1, $\hat{y}_1$ and $\hat{y}_2$ the point predictions of the regressors 1 and 2 respectively, and $\sigma_1^2$, $\sigma_2^2$ their per-prediction variances estimated from the quantile interval widths:

$$\sigma_k^2 = \left( \frac{\hat{Q}_{0.9}^{(k)} - \hat{Q}_{0.1}^{(k)}}{2 \times 1.28} \right)^2 \tag{4}$$

where 1.28 is the 90th percentile of the standard normal distribution. The total prediction variance of the gating model decomposes as:

$$\text{Var}(\hat{y}) = \underbrace{p\,\sigma_1^2 + (1-p)\,\sigma_2^2}_{\text{regressor uncertainty}} + \underbrace{p(1-p)\,(\hat{y}_1 - \hat{y}_2)^2}_{\text{gate uncertainty}} \tag{5}$$

The first term captures uncertainty intrinsic to each regressor, weighted by the gate probabilities. The second term captures the additional uncertainty arising when the classifier is uncertain which regime a node belongs to, and is maximised when $p = 0.5$ and the two regressors disagree. The square root $\sqrt{\text{Var}(\hat{y})}$ is reported in the same units as the prediction and used to characterise how model uncertainty varies across ancestral node ages.

### 4 Model accuracies for the phenotypes

| | Random | Family | Order | Class | Phylum | Domain (a-b) | Domain (b-a) |
|---|---|---|---|---|---|---|---|
| **Cross-validation** | | | | | | | |
| Balanced accuracy | $0.96 \pm 0.002$ | $0.96 \pm 0.004$ | $0.96 \pm 0.004$ | $0.94 \pm 0.01$ | $0.94 \pm 0.02$ | – | – |
| Recall | $0.96 \pm 0.002$ | $0.98 \pm 0.002$ | $0.97 \pm 0.007$ | $0.95 \pm 0.018$ | $0.95 \pm 0.02$ | – | – |
| F1 | $0.98 \pm 0.001$ | $0.97 \pm 0.002$ | $0.97 \pm 0.004$ | $0.96 \pm 0.01$ | $0.97 \pm 0.01$ | – | – |
| MCC | $0.94 \pm 0.004$ | $0.92 \pm 0.009$ | $0.92 \pm 0.009$ | $0.88 \pm 0.02$ | $0.89 \pm 0.04$ | – | – |
| ECE | $0.01 \pm 0.001$ | $0.02 \pm 0.002$ | $0.02 \pm 0.004$ | $0.04 \pm 0.008$ | $0.04 \pm 0.01$ | – | – |
| **Test** | | | | | | | |
| Balanced accuracy | $0.97 \pm 0.006$ | $0.96 \pm 0.01$ | $0.96 \pm 0.01$ | $0.92 \pm 0.04$ | $0.89 \pm 0.04$ | 0.85 | 0.96 |
| Recall | $0.98 \pm 0.004$ | $0.99 \pm 0.006$ | $0.98 \pm 0.01$ | $0.94 \pm 0.04$ | $0.95 \pm 0.02$ | 0.84 | 1.00 |
| F1 | $0.98 \pm 0.003$ | $0.98 \pm 0.008$ | $0.97 \pm 0.007$ | $0.95 \pm 0.01$ | $0.96 \pm 0.01$ | 0.88 | 0.93 |
| MCC | $0.94 \pm 0.004$ | $0.94 \pm 0.002$ | $0.93 \pm 0.02$ | $0.82 \pm 0.11$ | $0.82 \pm 0.17$ | 0.68 | 0.9 |
| ECE | $0.01 \pm 0.004$ | $0.02 \pm 0.007$ | $0.02 \pm 0.004$ | $0.05 \pm 0.02$ | $0.03 \pm 0.02$ | 0.05 | 0.04 |

Table 1: Oxygen use prediction performance across different taxonomic levels of train–test splits. Results are reported separately for cross-validation and held-out test evaluations. Values are shown as mean ± standard deviation across 30 splits at each taxonomic level.

| | Random | Family | Order | Class | Phylum |
|---|---|---|---|---|---|
| **Cross-validation** | | | | | |
| Balanced accuracy | $0.99 \pm 0.001$ | $0.99 \pm 0.001$ | $0.99 \pm 0.002$ | $0.98 \pm 0.007$ | $0.96 \pm 0.03$ |
| Recall | 1.00 | $0.99 \pm 0.001$ | $0.99 \pm 0.001$ | $0.98 \pm 0.02$ | $0.94 \pm 0.10$ |
| F1 | $0.99 \pm 0.001$ | $0.99 \pm 0.002$ | $0.98 \pm 0.004$ | $0.95 \pm 0.11$ | $0.93 \pm 0.06$ |
| MCC | $0.98 \pm 0.004$ | $0.99 \pm 0.004$ | $0.98 \pm 0.005$ | $0.97 \pm 0.01$ | $0.93 \pm 0.06$ |
| ECE | $0.008 \pm 0.001$ | $0.01 \pm 0.002$ | $0.01 \pm 0.002$ | $0.01 \pm 0.005$ | $0.04 \pm 0.02$ |
| **Test** | | | | | |
| Balanced accuracy | $0.99 \pm 0.005$ | $0.99 \pm 0.004$ | $0.99 \pm 0.005$ | $0.99 \pm 0.009$ | $0.97 \pm 0.02$ |
| Recall | 1.00 | 1.00 | 1.00 | $0.98 \pm 0.04$ | $0.97 \pm 0.05$ |
| F1 | $0.99 \pm 0.007$ | $0.99 \pm 0.003$ | $0.99 \pm 0.01$ | $0.97 \pm 0.02$ | $0.94 \pm 0.05$ |
| MCC | $0.99 \pm 0.01$ | $0.99 \pm 0.01$ | $0.98 \pm 0.01$ | $0.98 \pm 0.02$ | $0.93 \pm 0.06$ |
| ECE | $0.008 \pm 0.004$ | $0.007 \pm 0.004$ | $0.01 \pm 0.004$ | $0.01 \pm 0.009$ | $0.03 \pm 0.02$ |

Table 2: Cell envelope (mono/didermy) prediction performance across different taxonomic levels of train–test splits. Results are reported separately for cross-validation and held-out test evaluations. Values are shown as mean ± standard deviation across 30 splits at each taxonomic level.

| | Random | Family | Order | Class | Phylum |
|---|---|---|---|---|---|
| **Cross-validation** | | | | | |
| Balanced accuracy | $0.93 \pm 0.004$ | $0.89 \pm 0.01$ | $0.83 \pm 0.03$ | $0.67 \pm 0.05$ | $0.66 \pm 0.08$ |
| Recall | $0.88 \pm 0.008$ | $0.83 \pm 0.03$ | $0.81 \pm 0.05$ | $0.45 \pm 0.13$ | $0.34 \pm 0.18$ |
| F1 | $0.89 \pm 0.005$ | $0.85 \pm 0.02$ | $0.83 \pm 0.04$ | $0.51 \pm 0.42$ | $0.34 \pm 0.15$ |
| MCC | $0.87 \pm 0.007$ | $0.81 \pm 0.02$ | $0.80 \pm 0.05$ | $0.42 \pm 0.1$ | $0.34 \pm 0.15$ |
| ECE | $0.02 \pm 0.001$ | $0.03 \pm 0.004$ | $0.04 \pm 0.008$ | $0.09 \pm 0.03$ | $0.1 \pm 0.04$ |
| **Test** | | | | | |
| Balanced accuracy | $0.93 \pm 0.01$ | $0.9 \pm 0.02$ | $0.89 \pm 0.06$ | $0.71 \pm 0.12$ | $0.65 \pm 0.13$ |
| Recall | $0.88 \pm 0.03$ | $0.85 \pm 0.06$ | $0.80 \pm 0.14$ | $0.47 \pm 0.32$ | $0.43 \pm 0.28$ |
| F1 | $0.98 \pm 0.007$ | $0.86 \pm 0.05$ | $0.81 \pm 0.12$ | $0.53 \pm 0.30$ | $0.39 \pm 0.29$ |
| MCC | $0.88 \pm 0.02$ | $0.82 \pm 0.05$ | $0.80 \pm 0.11$ | $0.52 \pm 0.25$ | $0.39 \pm 0.25$ |
| ECE | $0.02 \pm 0.005$ | $0.03 \pm 0.01$ | $0.03 \pm 0.03$ | $0.07 \pm 0.06$ | $0.14 \pm 0.07$ |

Table 3: Sporulation prediction performance across different taxonomic levels of train–test splits. Results are reported separately for cross-validation and held-out test evaluations. Values are shown as mean ± standard deviation across 30 splits at each taxonomic level.

| | Random | Family | Order | Class | Phylum | Domain (a–b) | Domain (b–a) |
|---|---|---|---|---|---|---|---|
| **Cross-validation** | | | | | | | |
| Balanced accuracy | $0.91 \pm 0.005$ | $0.88 \pm 0.01$ | $0.88 \pm 0.01$ | $0.85 \pm 0.03$ | $0.85 \pm 0.02$ | – | – |
| Recall | $0.84 \pm 0.01$ | $0.8 \pm 0.02$ | $0.81 \pm 0.03$ | $0.75 \pm 0.06$ | $0.75 \pm 0.04$ | – | – |
| F1 | $0.88 \pm 0.008$ | $0.84 \pm 0.02$ | $0.82 \pm 0.02$ | $0.77 \pm 0.05$ | $0.78 \pm 0.04$ | – | – |
| MCC | $0.84 \pm 0.009$ | $0.81 \pm 0.02$ | $0.76 \pm 0.02$ | $0.73 \pm 0.04$ | $0.73 \pm 0.04$ | – | – |
| ECE | $0.02 \pm 0.001$ | $0.02 \pm 0.003$ | $0.03 \pm 0.006$ | $0.03 \pm 0.009$ | $0.04 \pm 0.01$ | – | – |
| RMSE | $5.55 \pm 0.06$ | $6.7 \pm 0.16$ | $7.14 \pm 0.3$ | $8.25 \pm 0.7$ | $8.39 \pm 0.8$ | – | – |
| $R^2$ | $0.85 \pm 0.004$ | $0.8 \pm 0.01$ | $0.77 \pm 0.02$ | $0.68 \pm 0.09$ | $0.71 \pm 0.04$ | – | – |
| **Test** | | | | | | | |
| Balanced accuracy | $0.91 \pm 0.01$ | $0.89 \pm 0.04$ | $0.88 \pm 0.05$ | $0.85 \pm 0.07$ | $0.82 \pm 0.08$ | 0.69 | 0.79 |
| Recall | $0.85 \pm 0.02$ | $0.8 \pm 0.08$ | $0.8 \pm 0.08$ | $0.76 \pm 0.13$ | $0.67 \pm 0.16$ | 0.6 | 0.88 |
| F1 | $0.89 \pm 0.01$ | $0.85 \pm 0.05$ | $0.82 \pm 0.09$ | $0.78 \pm 0.11$ | $0.73 \pm 0.13$ | 0.43 | 0.84 |
| MCC | $0.85 \pm 0.02$ | $0.81 \pm 0.06$ | $0.79 \pm 0.11$ | $0.73 \pm 0.11$ | $0.69 \pm 0.12$ | 0.31 | 0.59 |
| ECE | $0.02 \pm 0.006$ | $0.03 \pm 0.009$ | $0.03 \pm 0.02$ | $0.04 \pm 0.02$ | $0.04 \pm 0.02$ | 0.19 | 0.11 |
| RMSE | $5.44 \pm 0.21$ | $6.39 \pm 0.36$ | $6.85 \pm 0.6$ | $7.55 \pm 1.15$ | $7.7 \pm 0.96$ | 14.5 | 15.8 |
| $R^2$ | $0.86 \pm 0.01$ | $0.81 \pm 0.05$ | $0.74 \pm 0.11$ | $0.69 \pm 0.13$ | $0.64 \pm 0.16$ | $-0.35$ | 0.53 |

Table 4: Optimal growth temperature prediction performance across different taxonomic levels of train–test splits. Results are reported separately for cross-validation and held-out test evaluations. Values are shown as mean ± standard deviation across 30 splits at each taxonomic level.

| | Random | Family | Order | Class | Phylum | Domain (a–b) | Domain (b–a) |
|---|---|---|---|---|---|---|---|
| **Cross-validation** | | | | | | | |
| Balanced accuracy | $0.95 \pm 0.002$ | $0.86 \pm 0.01$ | $0.82 \pm 0.01$ | $0.71 \pm 0.03$ | $0.67 \pm 0.06$ | – | – |
| Recall | $0.95 \pm 0.002$ | $0.85 \pm 0.02$ | $0.84 \pm 0.02$ | $0.8 \pm 0.09$ | $0.69 \pm 0.13$ | – | – |
| F1 | $0.96 \pm 0.01$ | $0.87 \pm 0.01$ | $0.85 \pm 0.02$ | $0.78 \pm 0.06$ | $0.7 \pm 0.11$ | – | – |
| MCC | $0.90 \pm 0.04$ | $0.72 \pm 0.02$ | $0.63 \pm 0.03$ | $0.44 \pm 0.08$ | $0.35 \pm 0.13$ | – | – |
| ECE | $0.07 \pm 0.008$ | $0.14 \pm 0.04$ | $0.18 \pm 0.07$ | $0.25 \pm 0.12$ | $0.26 \pm 0.1$ | – | – |
| RMSE | $4 \pm 0.05$ | $6.67 \pm 0.23$ | $7.73 \pm 0.42$ | $9.7 \pm 0.52$ | $11.1 \pm 1.39$ | – | – |
| $R^2$ | $0.88 \pm 0.03$ | $0.73 \pm 0.02$ | $0.64 \pm 0.02$ | $0.4 \pm 0.08$ | $0.23 \pm 0.15$ | – | – |
| **Test** | | | | | | | |
| Balanced accuracy | $0.95 \pm 0.005$ | $0.85 \pm 0.05$ | $0.81 \pm 0.05$ | $0.72 \pm 0.11$ | $0.69 \pm 0.06$ | 0.53 | 0.56 |
| Recall | $0.95 \pm 0.006$ | $0.84 \pm 0.07$ | $0.83 \pm 0.08$ | $0.77 \pm 0.21$ | $0.76 \pm 0.06$ | 0.83 | 0.58 |
| F1 | $0.96 \pm 0.004$ | $0.87 \pm 0.06$ | $0.83 \pm 0.07$ | $0.71 \pm 0.24$ | $0.72 \pm 0.17$ | 0.68 | 0.56 |
| MCC | $0.9 \pm 0.001$ | $0.71 \pm 0.09$ | $0.64 \pm 0.11$ | $0.44 \pm 0.21$ | $0.41 \pm 0.11$ | 0.57 | 0.13 |
| ECE | $0.07 \pm 0.007$ | $0.12 \pm 0.03$ | $0.14 \pm 0.04$ | $0.18 \pm 0.08$ | $0.21 \pm 0.11$ | 0.26 | 0.35 |
| RMSE | $3.9 \pm 0.16$ | $6.45 \pm 0.88$ | $7.21 \pm 0.91$ | $8.74 \pm 1.72$ | $9.7 \pm 1.51$ | 13.01 | 10.94 |
| $R^2$ | $0.88 \pm 0.003$ | $0.72 \pm 0.08$ | $0.61 \pm 0.09$ | $0.22 \pm 0.56$ | $0.24 \pm 0.24$ | $-0.01$ | $-0.02$ |

Table 5: GC content prediction performance across different taxonomic levels of train–test splits. Results are reported separately for cross-validation and held-out test evaluations. Values are shown as mean ± standard deviation across 30 splits at each taxonomic level.

#### 5 Markov Blanket algorithm

We use the Incremental Association Markov blanket (IAMB) algorithm to find the Markov blankets (MB) for a given target variable $T$, and the feature set $V$ [**?**, **?**]. The algorithm is outlined below.

We initialize

$$\mathrm{MB}(T) = \emptyset.$$

**Forward phase.** In the forward phase, features are iteratively added to $\mathrm{MB}(T)$ by selecting the variable that maximizes the conditional mutual information with the target given the current Markov blanket.

At each iteration, we select

$$X_{\max} = \arg \max_{X \in V \setminus (\mathrm{MB}(T) \cup \{T\})} \mathrm{CMI}(X; T \mid \mathrm{MB}(T)),$$

where $\mathrm{CMI}(X; Y \mid Z)$ is the conditional mutual information between random variables $X$ and $Y$ given $Z$, and is defined as

$$\mathrm{CMI}(X; Y \mid Z) = \sum_{x,y,z} p(x, y, z) \log \frac{p(x, y \mid z)}{p(x \mid z)\, p(y \mid z)}, \tag{6}$$

where $p(x, y, z)$ denotes the joint probability distribution of $(X, Y, Z)$, and $p(\cdot \mid z)$ are the corresponding conditional distributions.

If

$$X_{\max} \not\perp\!\!\!\perp T \mid \mathrm{MB}(T),$$

we update

$$\mathrm{MB}(T) \leftarrow \mathrm{MB}(T) \cup \{X_{\max}\}.$$

This procedure is repeated until $\mathrm{MB}(T)$ no longer changes.

**Backward phase.** In the backward phase, spurious features are removed from $\mathrm{MB}(T)$ by testing conditional independence with the target given the remaining blanket variables.

For each $X \in \mathrm{MB}(T)$, we test whether

$$X \perp\!\!\!\perp T \mid \mathrm{MB}(T) \setminus \{X\}.$$

If this condition holds, we update

$$\mathrm{MB}(T) \leftarrow \mathrm{MB}(T) \setminus \{X\}.$$

The final set $\mathrm{MB}(T)$ is returned as the estimated Markov blanket of the target variable $T$.

### 6 Model performance on top-SHAP features and Markov blankets for all phenotypes

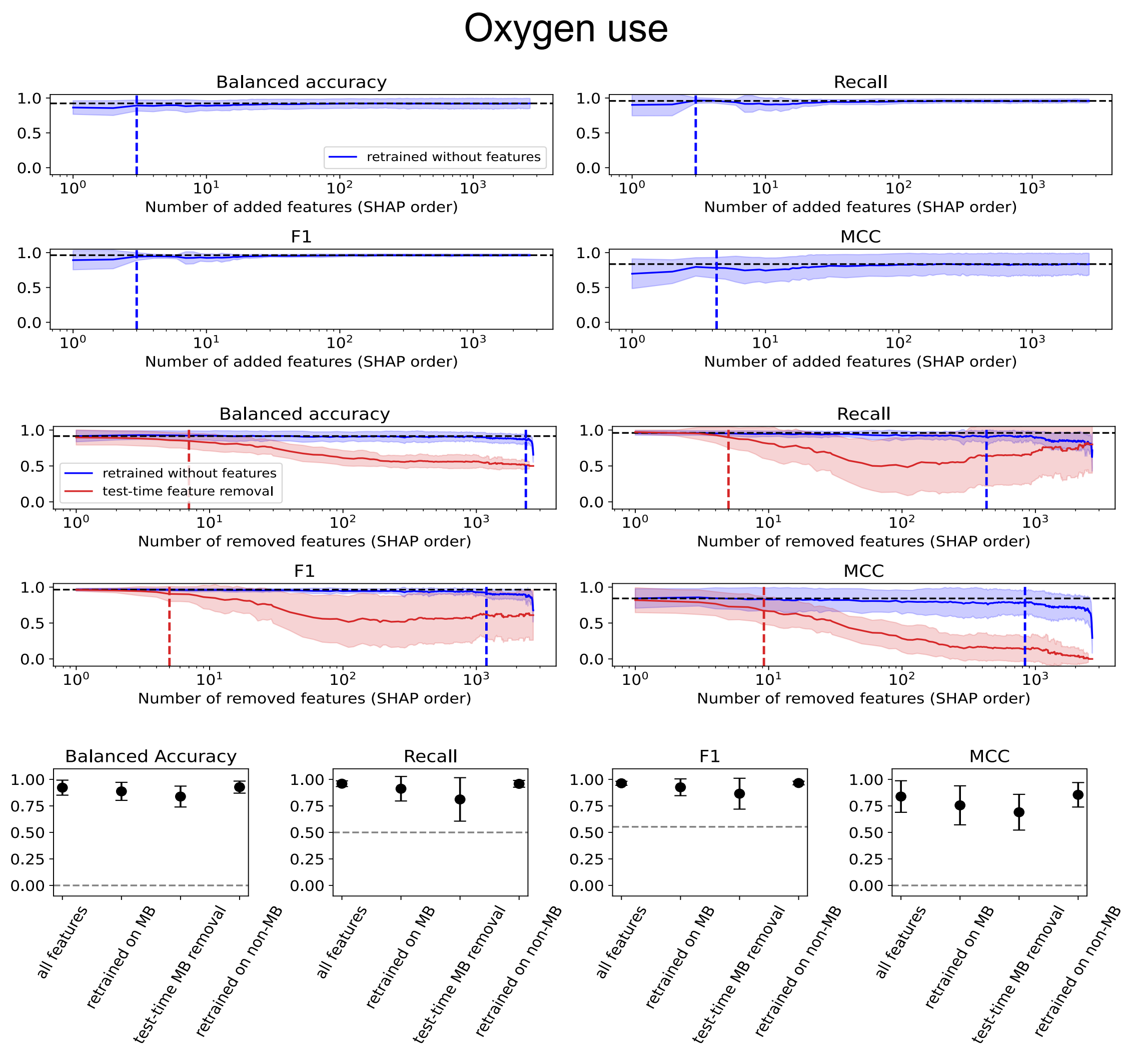


Figure 4: Oxygen use predictor performance under phylum-level test holdout using selected features. Top: Top-SHAP feature addition and removal experiments. For feature removal, we evaluate two settings: removal during both training and testing, and removal at test time only. Bottom: Model accuracies obtained using Markov blanket and non-Markov blanket features, compared with the full feature set.

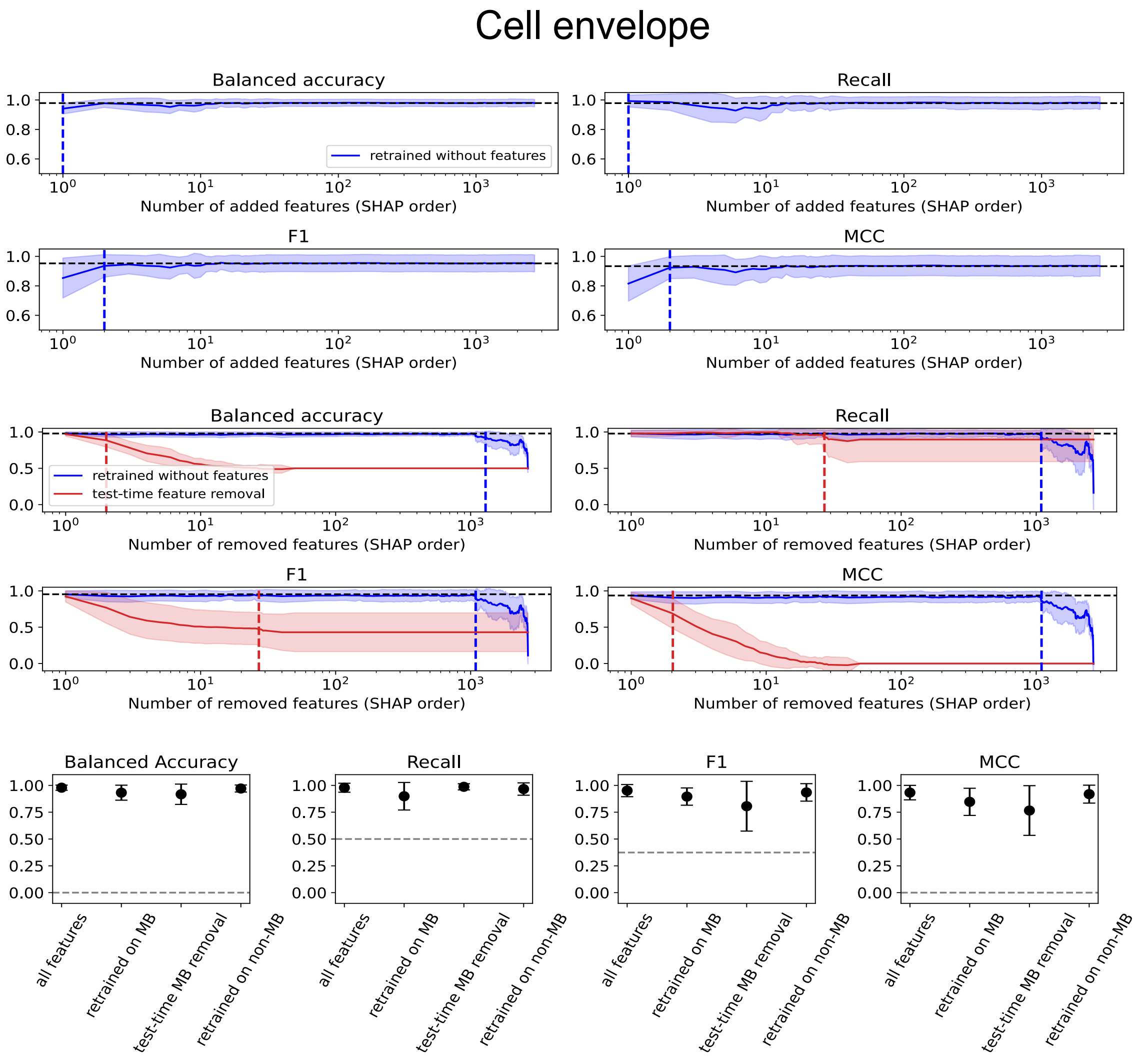


Figure 5: Cell envelope predictor performance under phylum-level test holdout using selected features. Top: Top-SHAP feature addition and removal experiments. For feature removal, we evaluate two settings: removal during both training and testing, and removal at test time only. Bottom: Model accuracies obtained using Markov blanket and non-Markov blanket features, compared with the full feature set.

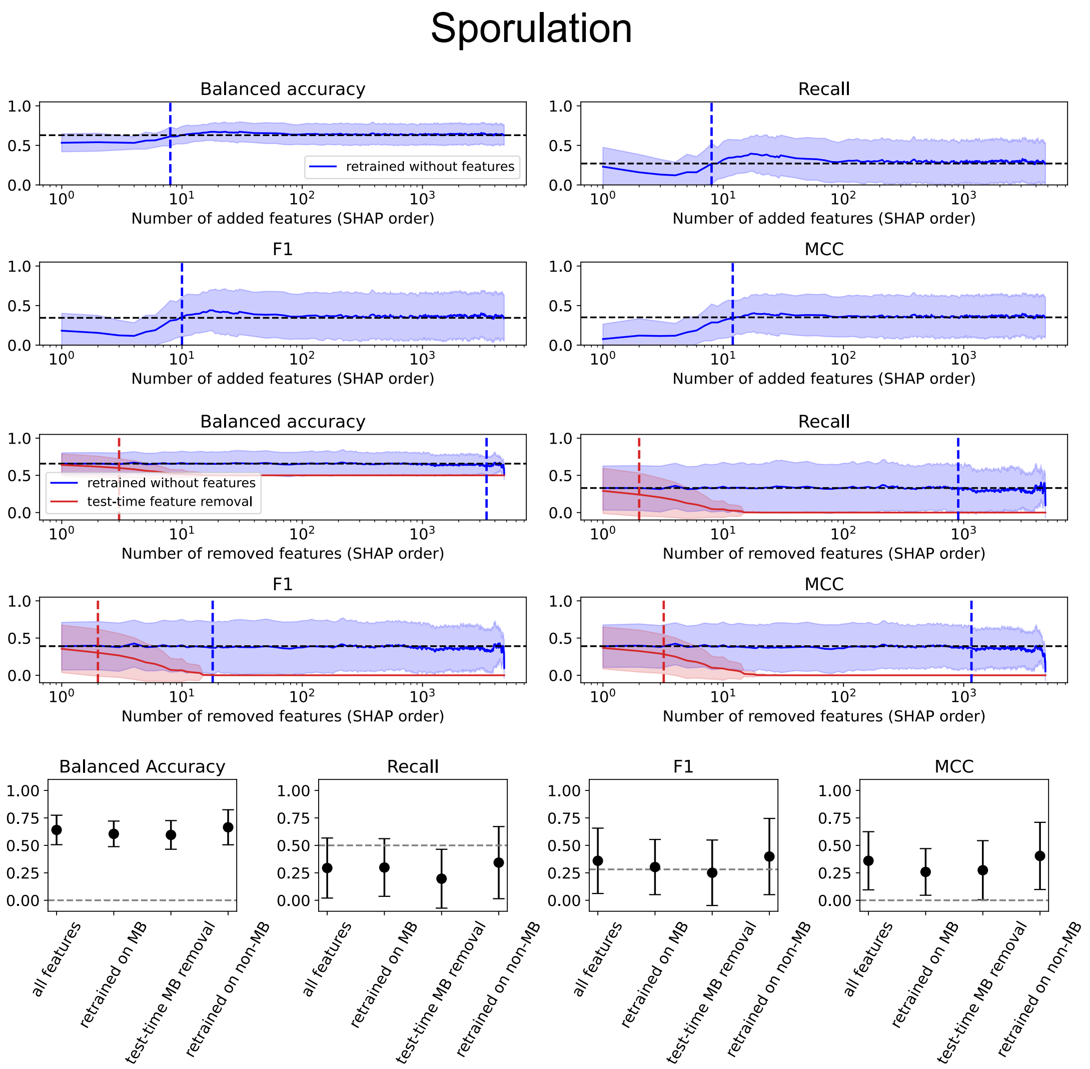


Figure 6: Sporulation predictor performance under phylum-level test holdout using selected features. Top: Top-SHAP feature addition and removal experiments. For feature removal, we evaluate two settings: removal during both training and testing, and removal at test time only. Bottom: Model accuracies obtained using Markov blanket and non-Markov blanket features, compared with the full feature set.

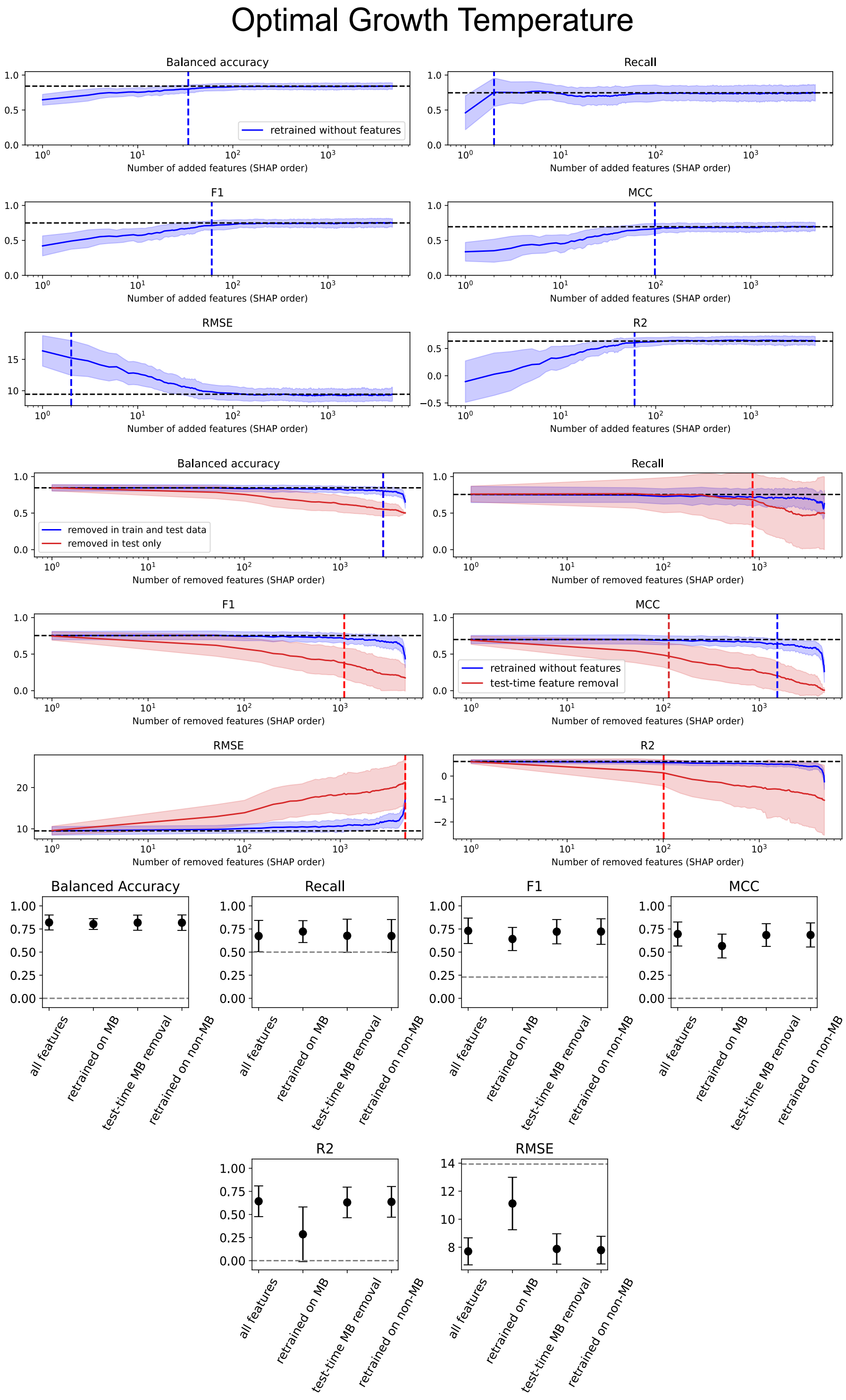


Figure 7: OGT predictor performance under phylum-level test holdout using selected features. Top: Top-SHAP feature addition and removal experiments. For feature removal, we evaluate two settings: removal during both training and testing, and removal at test time only. Bottom: Model accuracies obtained using Markov blanket and non-Markov blanket features, compared with the full feature set.

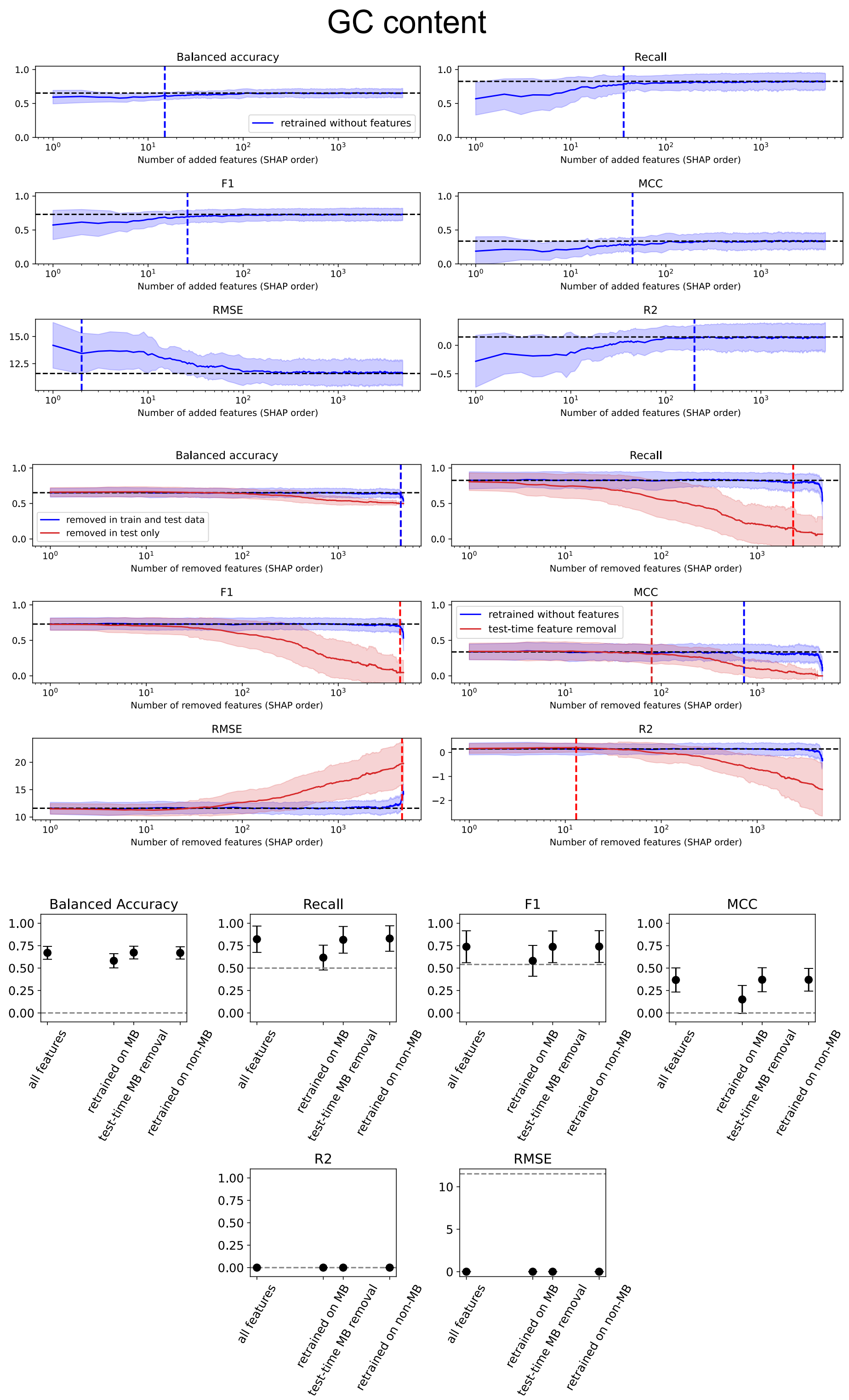


Figure 8: GC content predictor performance under phylum-level test holdout using selected features. Top: Top-SHAP feature addition and removal experiments. For feature removal, we evaluate two settings: removal during both training and testing, and removal at test time only. Bottom: Model accuracies obtained using Markov blanket and non-Markov blanket features, compared with the full feature set.

### 7 Top-SHAP features and Markov blankets for cell envelope, sporulation, OGT, and GC content

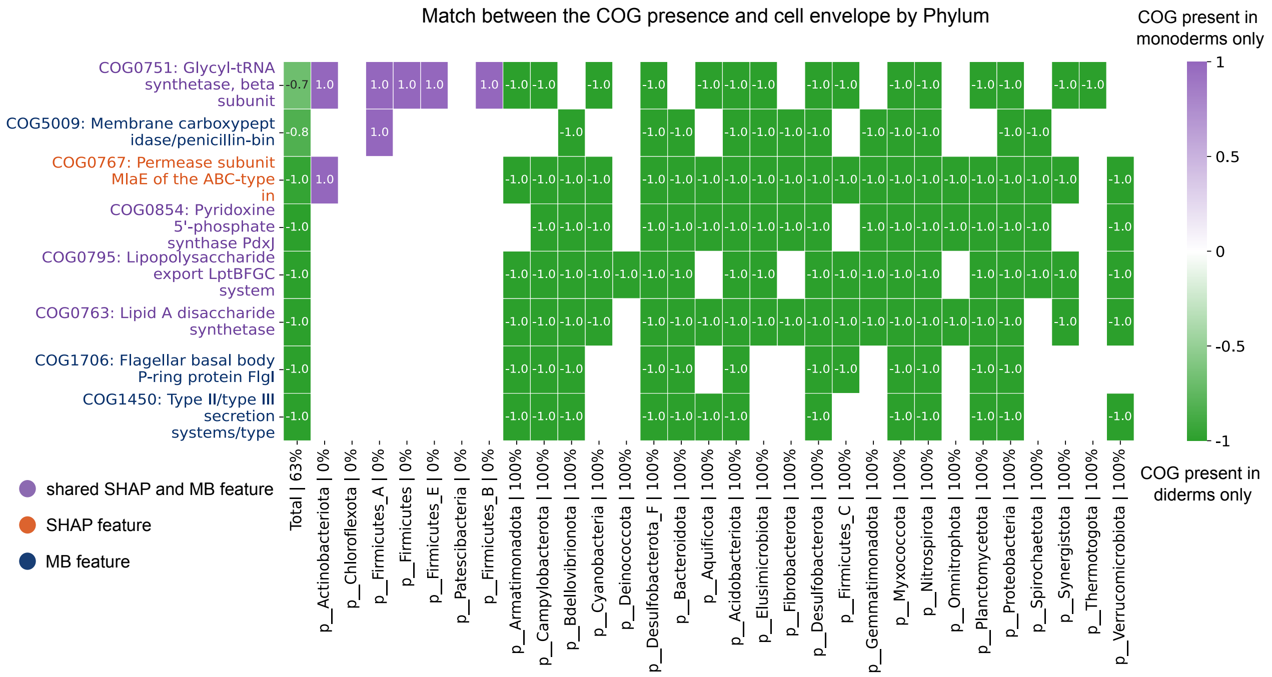


Figure 9: Top SHAP-ranked features and Markov blanket (MB) features associated with cell envelope phenotype, and their presence–phenotype pattern overlap across phyla. Feature colors indicate membership in the top-SHAP set (orange), the MB (dark blue), or their intersection (purple). The score quantifies the overlap between COG presence and phenotype occurrence (see Methods, ??). A score of +1 (purple) indicates perfect agreement between COG presence and monodermy, whereas a score of -1 (green) indicates perfect agreement between COG presence and didermy, i.e. the COG is present exclusively in diderm organisms.

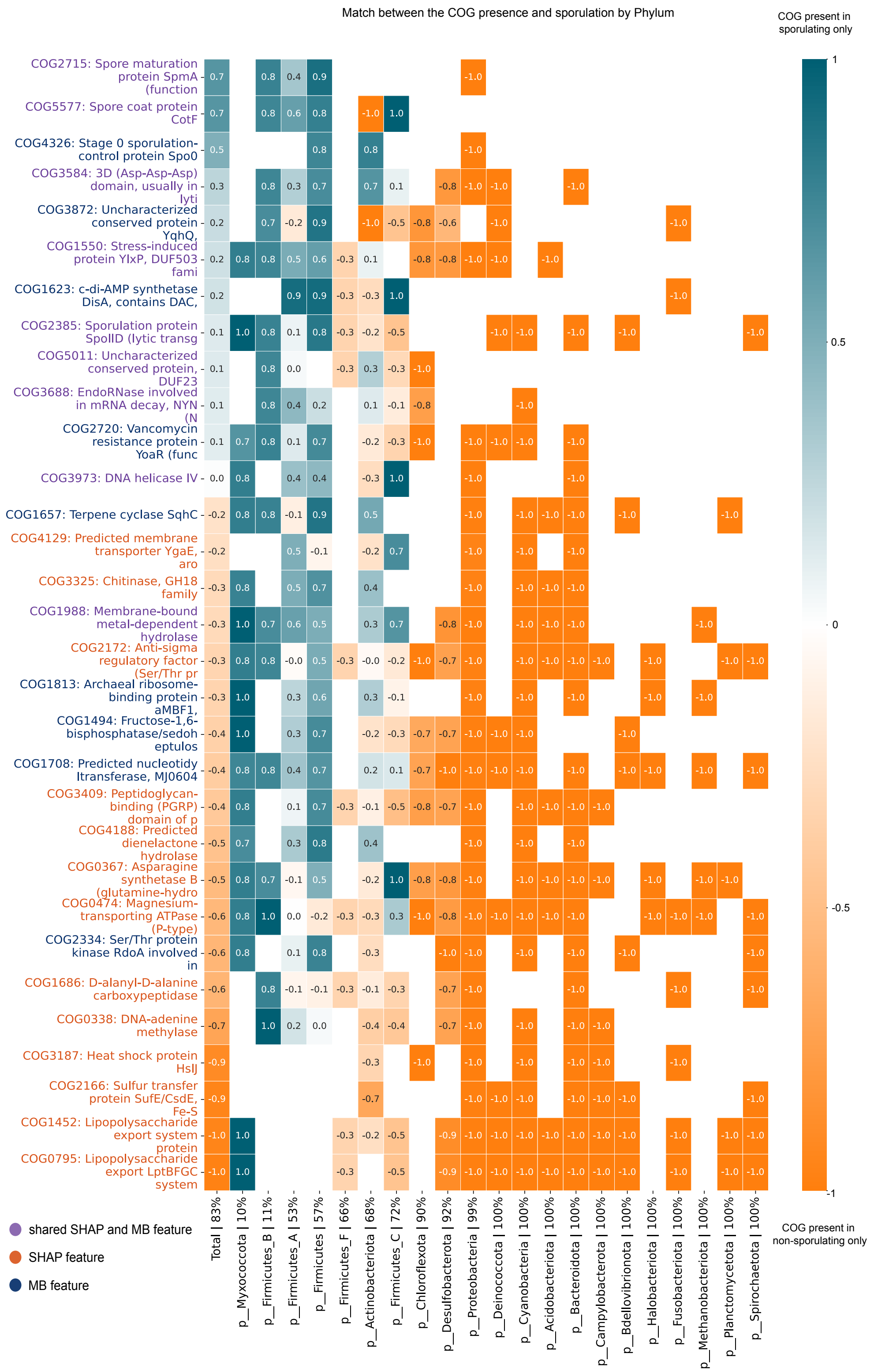


Figure 10: Top SHAP-ranked features and Markov blanket (MB) features associated with sporulation phenotype, and their presence–phenotype pattern overlap across phyla. Feature colors indicate membership in the top-SHAP set (orange), the MB (dark blue), or their intersection (purple). The score quantifies the overlap between COG presence and phenotype occurrence (see Methods). A score of +1 (teal) indicates perfect agreement between COG presence and sporulation, whereas a score of -1 (orange) indicates perfect agreement between COG presence and non-sporulation, i.e. the COG is present exclusively in non-sporulating organisms.

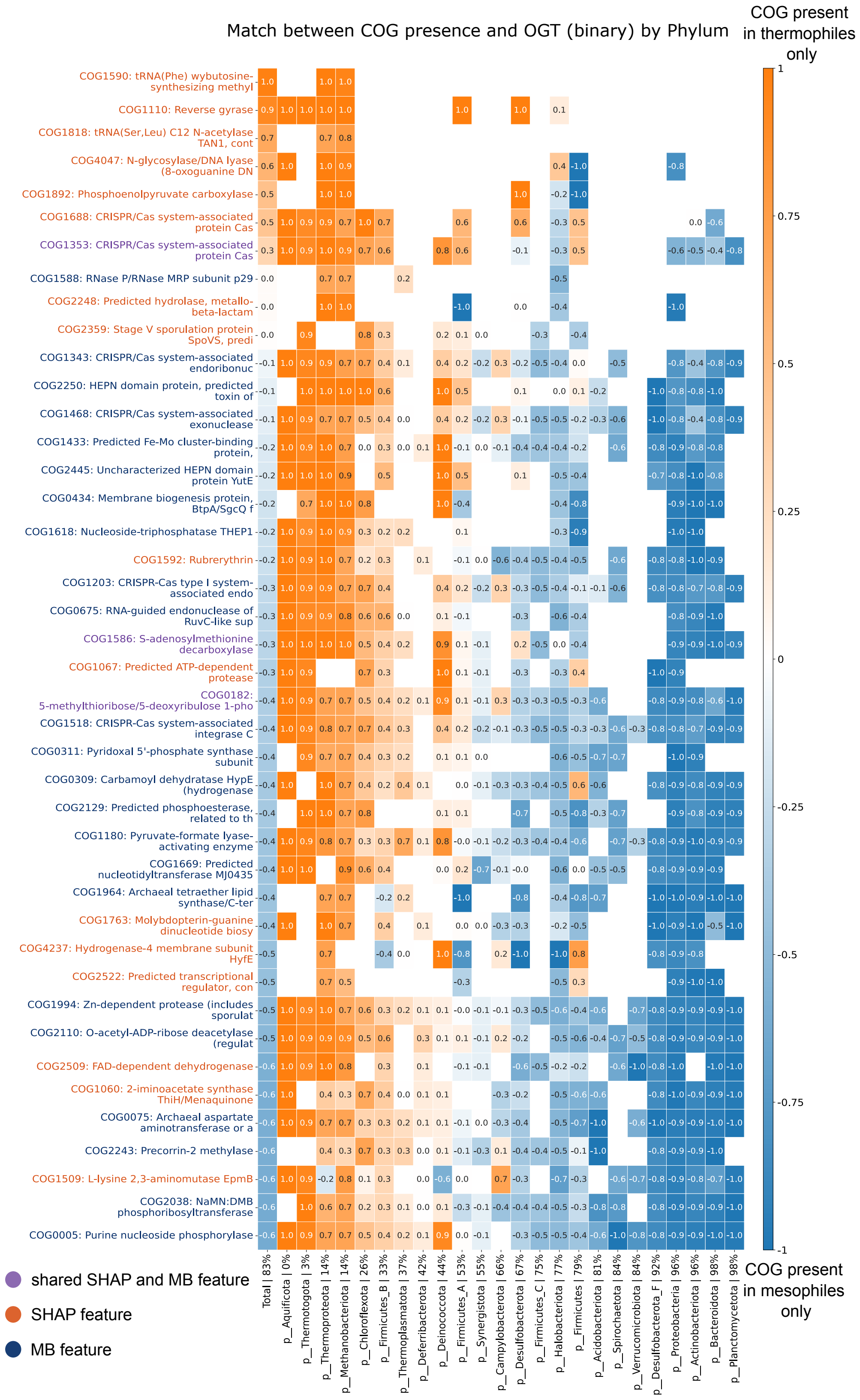


Figure 11: Top SHAP-ranked features and Markov blanket (MB) features associated with optimal growth temperature (OGT) phenotype, and their presence–phenotype pattern overlap across phyla (Part 1). Feature colors indicate membership in the top-SHAP set (orange), the MB (dark blue), or their intersection (purple). The score quantifies the overlap between COG presence and phenotype occurrence (see Methods, ??). A score of +1 (orange) indicates perfect agreement between COG presence and thermophiles, whereas a score of -1 (blue) indicates perfect agreement between COG presence and mesophiles, i.e. the COG is present exclusively in mesophile organisms.

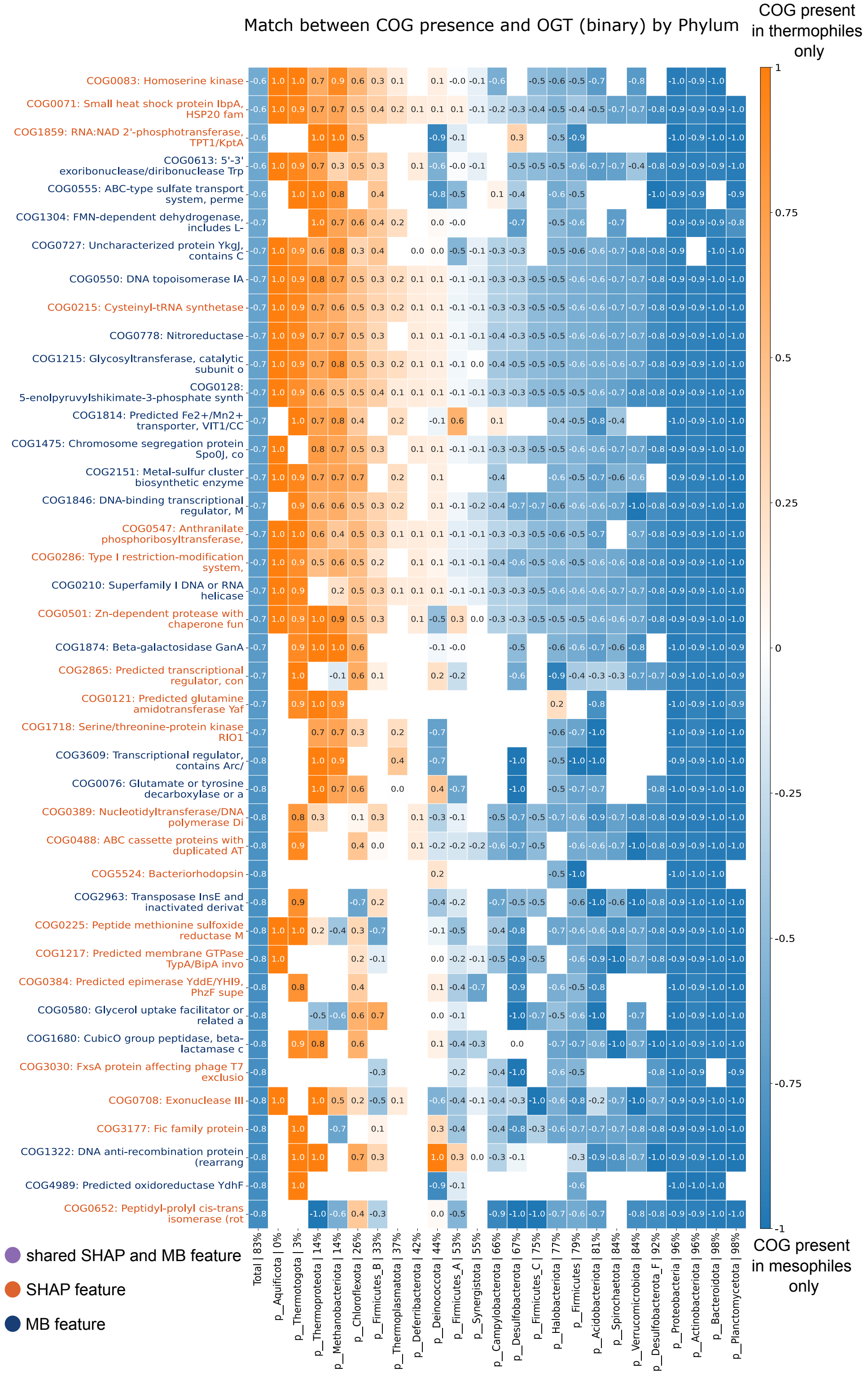


Figure 12: Top SHAP-ranked features and Markov blanket (MB) features associated with optimal growth temperature (OGT) phenotype, and their presence–phenotype pattern overlap across phyla, Part 2 of Fig. 11.

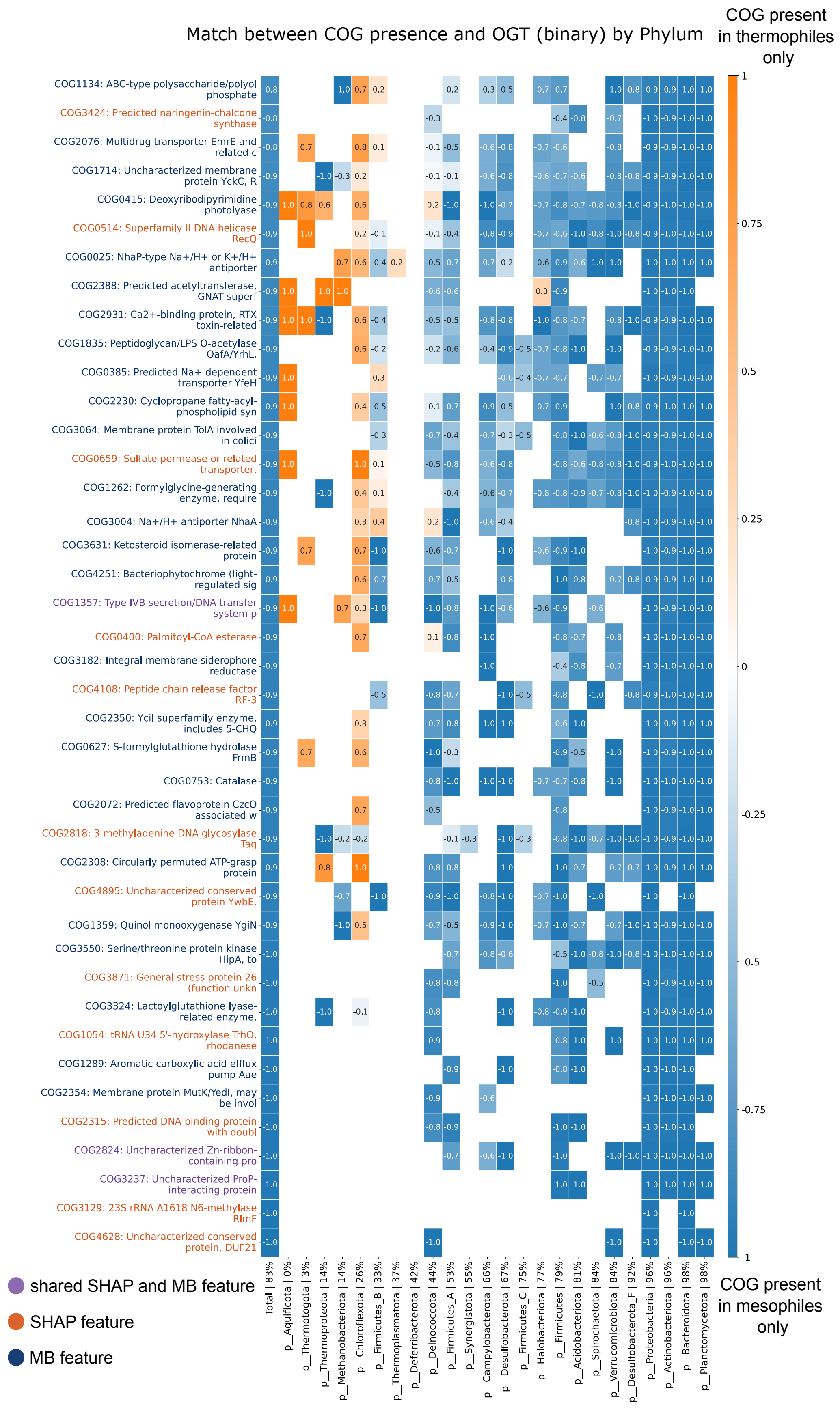


Figure 13: Top SHAP-ranked features and Markov blanket (MB) features associated with optimal growth temperature (OGT) phenotype, and their presence–phenotype pattern overlap across phyla, Part 3 of Fig. 11.

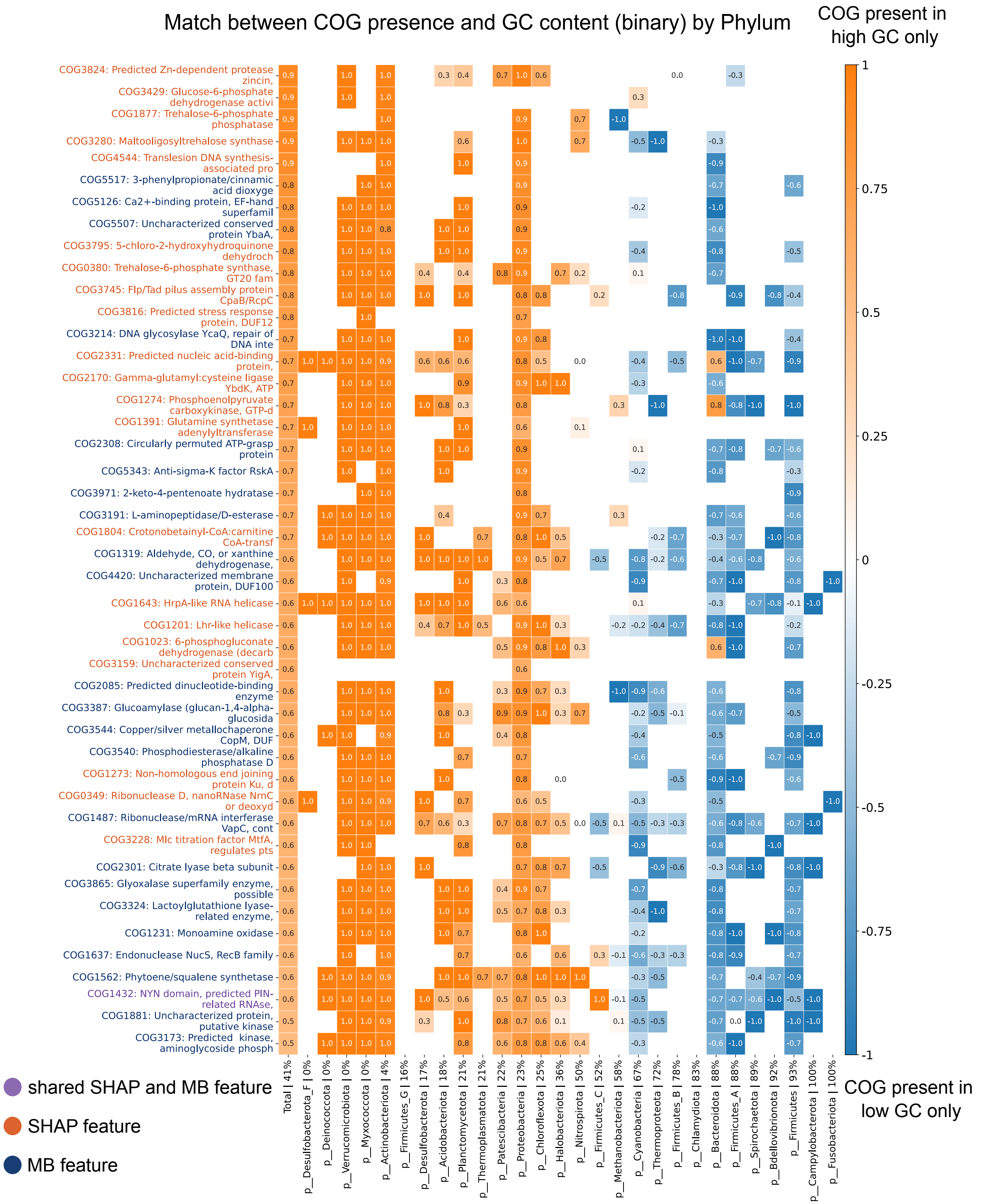


Figure 14: Top SHAP-ranked features and Markov blanket (MB) features associated with GC content phenotype, and their presence–phenotype pattern overlap across phyla (Part 1). Feature colors indicate membership in the top-SHAP set (orange), the MB (dark blue), or their intersection (purple). The score quantifies the overlap between COG presence and phenotype occurrence (see Methods). A score of +1 (orange) indicates perfect agreement between COG presence and high GC, whereas a score of -1 (blue) indicates perfect agreement between COG presence and low GC, i.e. the COG is present exclusively in low GC organisms.

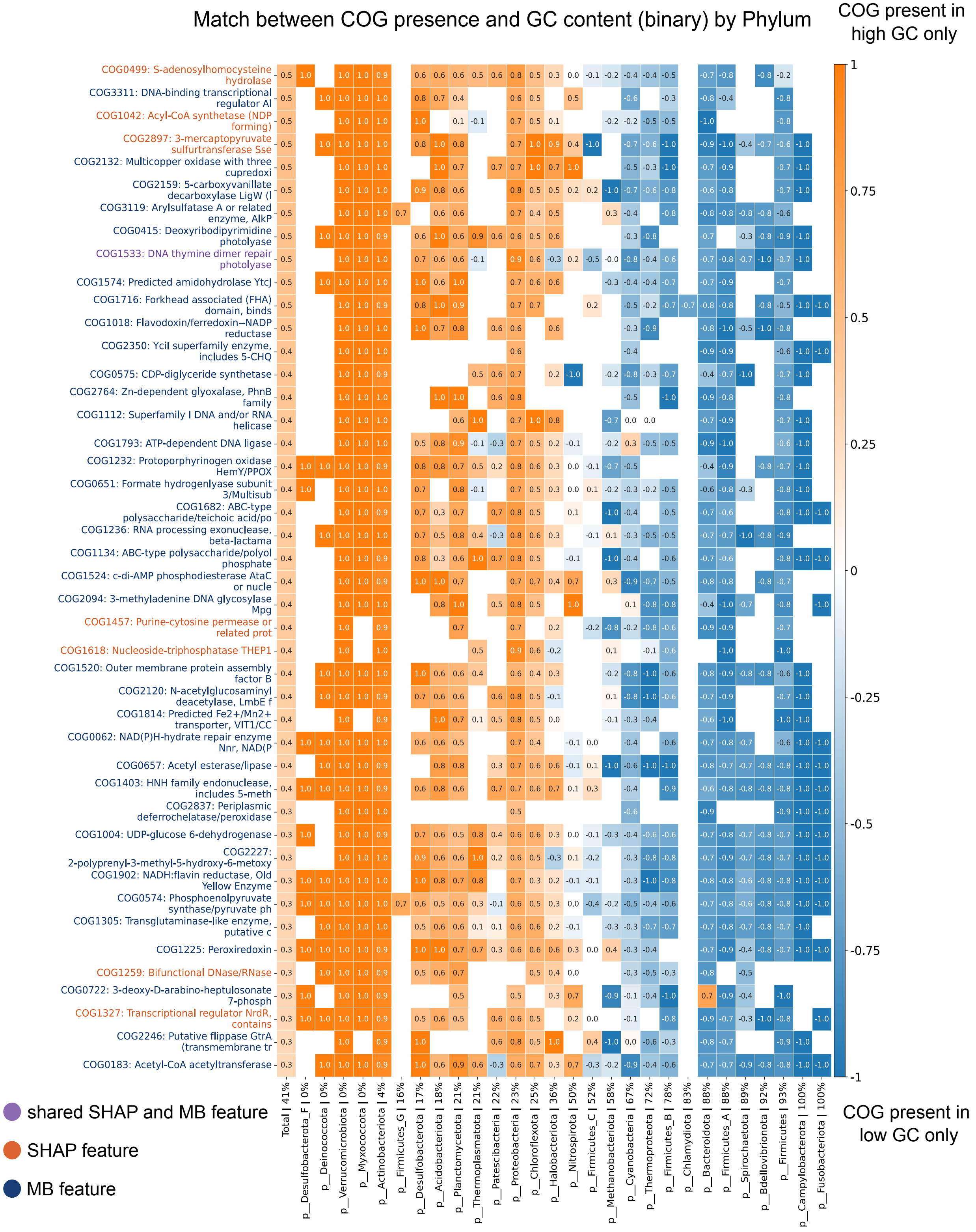


Figure 15: Top SHAP-ranked features and Markov blanket (MB) features associated with GC content phenotype, and their presence–phenotype pattern overlap across phyla. Part 2 of Fig. 14.

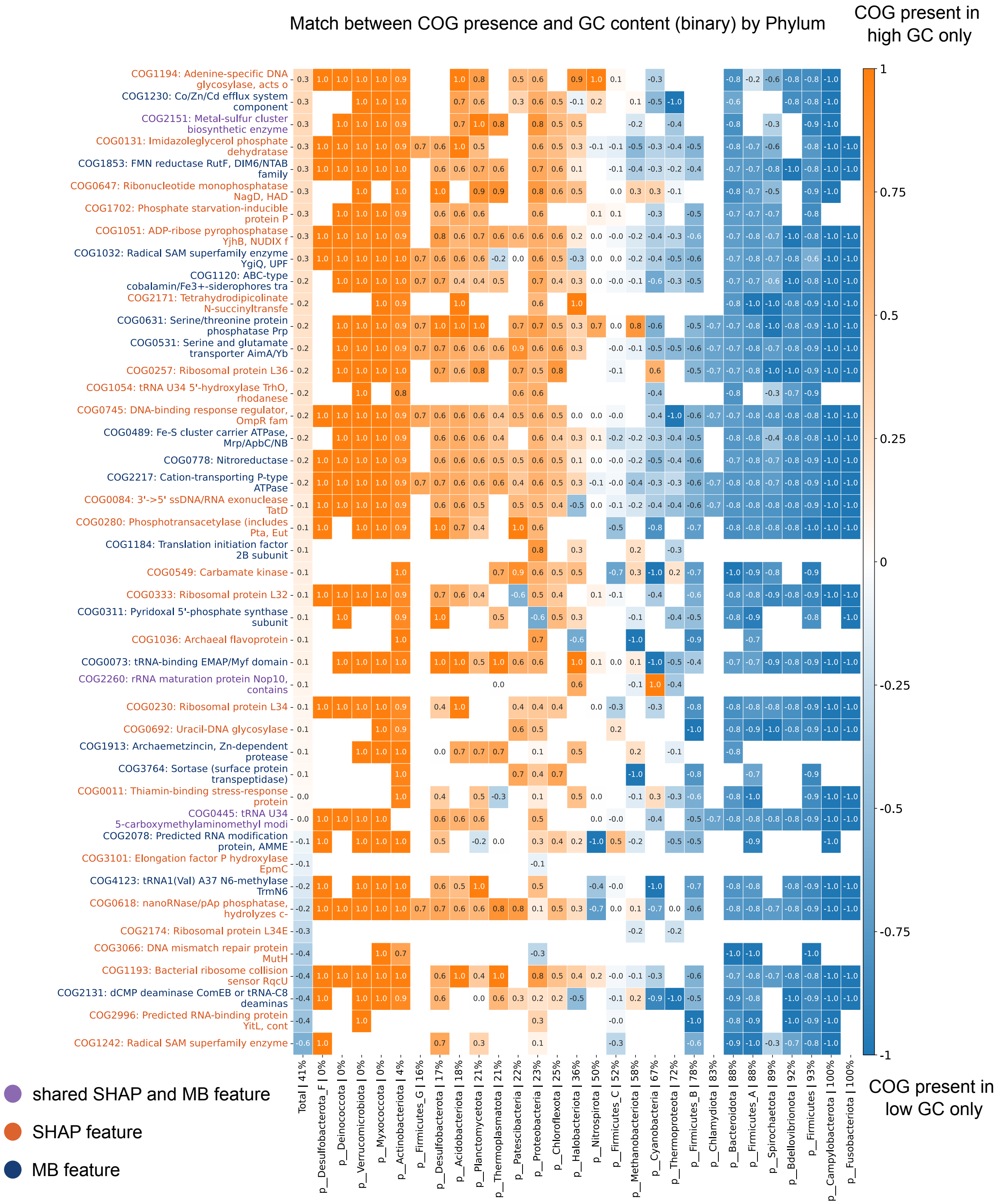


Figure 16: Top SHAP-ranked features and Markov blanket (MB) features associated with GC content phenotype, and their presence–phenotype pattern overlap across phyla. Part 3 of Fig. 14.

### 8 Training noise-robust models

#### 8.1 Train data augmentation

Training data are augmented with noisy genome copies, in which false positive and false negative perturbations are introduced according to Eqs. **??**. Specifically, the false positive and false negative rates, $s_{FP}$ and $s_{FN}$, are independently sampled from uniform, exponential, or gamma distributions. Each distribution is parameterized by its mean value, denoted $\lambda_{FP}$ and $\lambda_{FN}$, respectively. For the gamma distributions, the scale parameter is fixed to 0.5. We consider $\lambda_{FN} \in \{0, 0.2, 0.5, 1, 2, 4\}$ and $\lambda_{FP} \in \{0, 0.05, 0.1, 0.15, 0.2, 0.5\}$. Representative examples of these distributions are shown in Fig. 17.

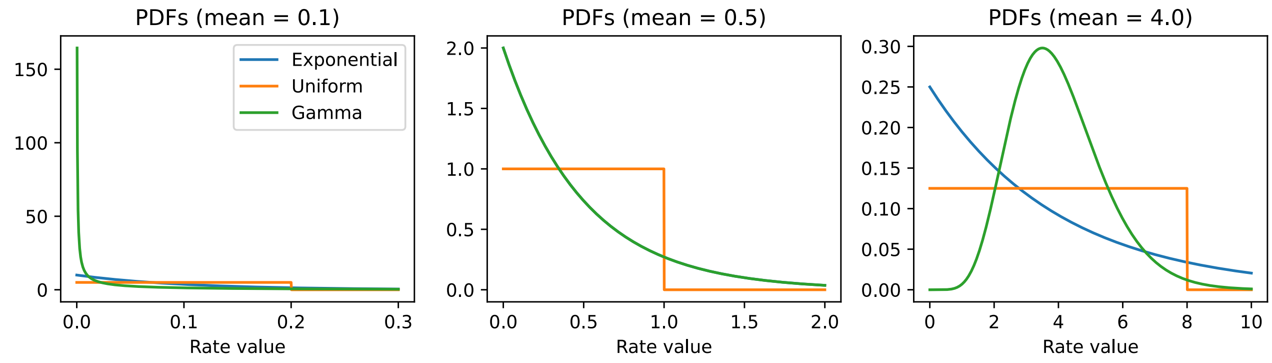


Figure 17: Exponential, Uniform, and Gamma distributions for the false positive and false negative rate sampling. The three examples demonstrate the three groups of the distributions for different mean values. The scale parameter in the gamma distributions is fixed to 0.5

The choice of the $\lambda_{FN}$ and $\lambda_{FP}$ parameters determines the level of noise injected into the augmented training dataset. In Fig. 18, we illustrate three representative noise regimes using 2-dimensional t-SNE projections of the augmented training data. When the noise level is too low, the noisy samples remain very close to the original genomes and therefore introduce little additional variation into the dataset, see Fig. 18, left. In contrast, when the noise level is too high, the augmented genomes lose much of their meaningful biological signal, resulting in a strongly distorted data structure, see Fig. 18, right. Our goal is to identify an intermediate noise regime that improves model generalization while preserving the overall structure of the augmented training data, see Fig. 18, center.

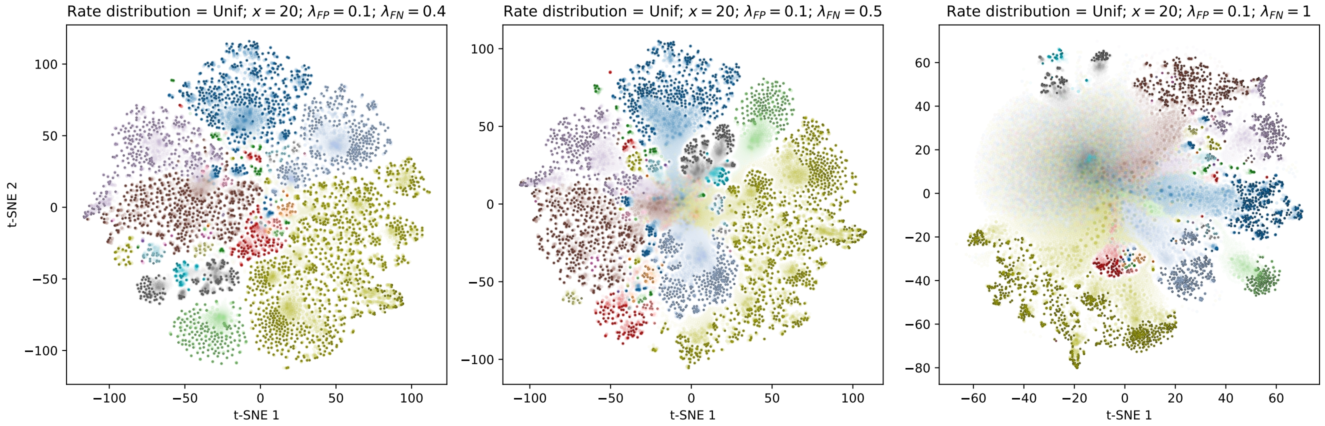


Figure 18: 2-dimensional t-SNE projections of the noise-augmented oxygen-use dataset for $x = 20$ noisy genome copies per original genome. False positive and false negative rates are sampled from uniform distributions with $\lambda_{FP} = 0.1$ and $\lambda_{FN} = 0.4$ (left), $\lambda_{FP} = 0.1$ and $\lambda_{FN} = 0.5$ (center), and $\lambda_{FP} = 0.1$ and $\lambda_{FN} = 1.0$ (right). Different colors correspond to different phyla. Bright colors represent the original genomes, whereas semi-transparent points represent the noisy augmented genomes.

#### 8.2 Model performance

We systematically explore model performance across different noise augmentation regimes defined by the parameters $(x, \lambda_{FN}, \lambda_{FP})$, where $x$ is the number of noisy copies generated per original genome, and $\lambda_{FN}$ and $\lambda_{FP}$ are the mean parameters of the distributions used to sample the false negative and false positive rates $r_{FN}$ and $r_{FP}$, respectively. We consider three sampling distributions (Exponential, Uniform, and Gamma), as defined in Section 8.1.

Our goal is to identify, for each phenotype, the noise augmentation regime that maximises model performance across a range of test-time noise levels. To prevent data leakage in the selection of $(x, \lambda_{FN}, \lambda_{FP})$ and rate distributions, we partition the annotation datasets into training, test, and validation sets in a $60\%/20\%/20\%$ ratio using phylum-level taxonomic holdout. The test set is used to evaluate model performance under different noise augmentation regimes and select the optimal configuration; the validation set is held out entirely and used only to report final model performance, providing an unbiased estimate of generalisation.

For each noise configuration, we augment the training data and train XGBoost models across multiple train/test/validation splits. Each trained model is then evaluated on noisy test datasets across a range of test-time false negative and false positive noise rates, denoted $r_{FN}$ and $r_{FP}$ respectively. Model performance is assessed using three complementary metrics: MCC, $F_1$, and the ECE score.

We define the MCC, $F_1$, and ECE score performance surfaces of a model across the test-time noise space as $M(r_{FN}, r_{FP})$, $F(r_{FN}, r_{FP})$, and $B(r_{FN}, r_{FP})$, respectively. We then integrate these surfaces over a selected noise range $r_{FN} \in [0.4, 0.8]$ and $r_{FP} \in [0.05, 0.15]$. This range was chosen to emphasise medium-to-high noise levels, reflecting the expected phenotype prediction quality of noisy reconstructed ancestral genomes. The overall robustness of a model is then quantified by the integrals

$$S_{\text{MCC}} = \int_{0.05}^{0.15} \int_{0.4}^{0.8} M(r_{FN}, r_{FP})\, dr_{FN}\, dr_{FP}, \tag{7}$$

$$S_{\text{F1}} = \int_{0.05}^{0.15} \int_{0.4}^{0.8} F(r_{FN}, r_{FP})\, dr_{FN}\, dr_{FP}, \tag{8}$$

$$S_{\text{ECE}} = \int_{0.05}^{0.15} \int_{0.4}^{0.8} B(r_{FN}, r_{FP})\, dr_{FN}\, dr_{FP}. \tag{9}$$

We normalise Eqs. (7), (8), (9) by the area of the integration domain, given by $\int_{0.05}^{0.15} \int_{0.4}^{0.8} dr_{FP}\, dr_{FN} = 0.04$, such that the resulting summary statistics represent the average model performance over the selected noise range rather than the cumulative sum. This yields the normalised integral scores $\bar{S}_{\text{MCC}}$, $\bar{S}_{F_1}$, and $\bar{S}_{\text{ECE}}$, which are shown in Fig. 22 for the oxygen use phenotype. The corresponding figures for the remaining phenotypes are provided in the Supplementary Materials.

The optimal noise augmentation regime corresponds to models achieving the highest $\bar{S}_{\text{MCC}}$ and $\bar{S}_{F_1}$, and the lowest $\bar{S}_{\text{ECE}}$. Since no single model simultaneously maximizes all three criteria across all noise augmentation regimes, model selection is performed with respect to the Pareto front - the set of models for which no other configuration achieves a strictly better performance on all three metrics simultaneously. The position of a model on the Pareto front reflects the optimal level of noise augmentation that improves generalization to noisy test data without destroying the phenotypic signal in the training data.

The noise augmentation regimes selected for each phenotype, used both to demonstrate performance improvement on the validation dataset acros different noise ranges and to train the final model for ancestral phenotype prediction, are reported in Table 6. The improved model performance on the validation data is shown in Fig. **??** in the main text.

| | Rate distribution | $\lambda_{FP}$ | $\lambda_{FN}$ | $x$ |
|---|---|---|---|---|
| Oxygen use | Uniform | 0.1 | 0.5 | 100 |
| Cell envelope | Exponential | 0.1 | 0.5 | 100 |
| Sporulation | Uniform | 0.1 | 1.0 | 100 |
| OGT | Exponential | 0.1 | 0.5 | 100 |
| GC content | Exponential | 0.1 | 1.0 | 100 |

Table 6: Noise regimes selected for each phenotype to train the final model used for ancestral phenotype prediction.

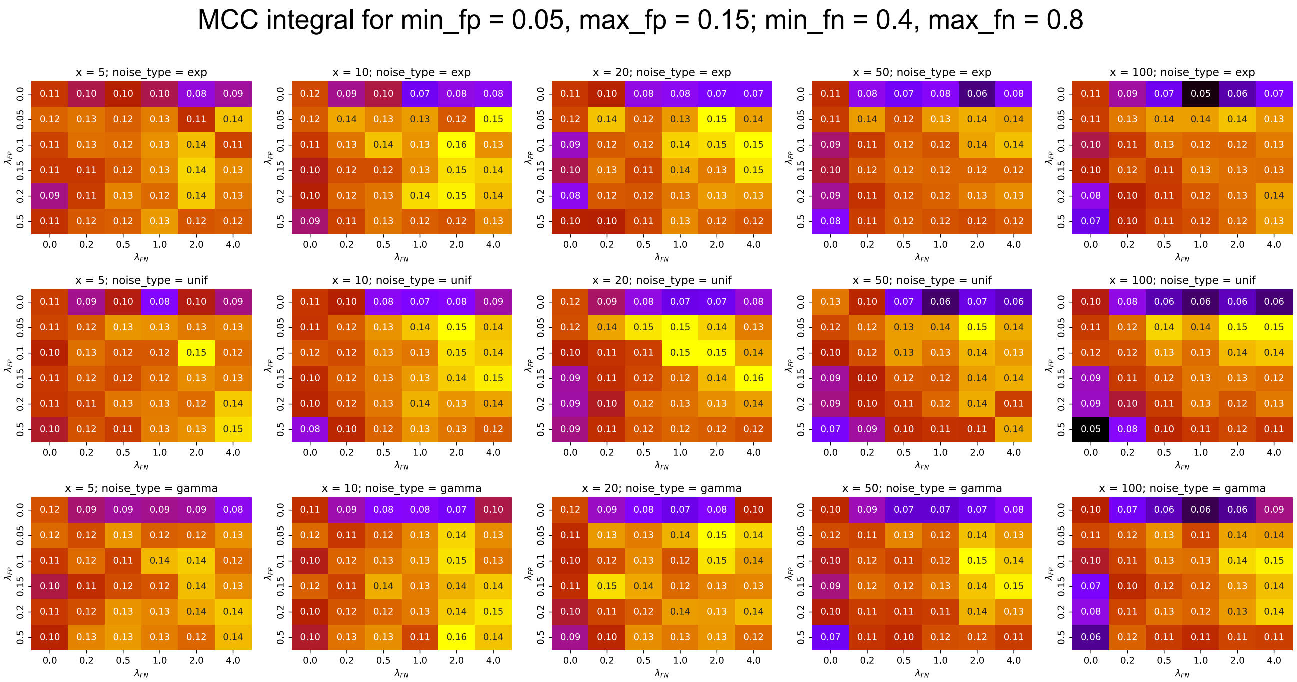


Figure 19: Normalised integral scores $\bar{S}_{\mathrm{MCC}}$ across different noise augmentation regimes for the oxygen use phenotype.

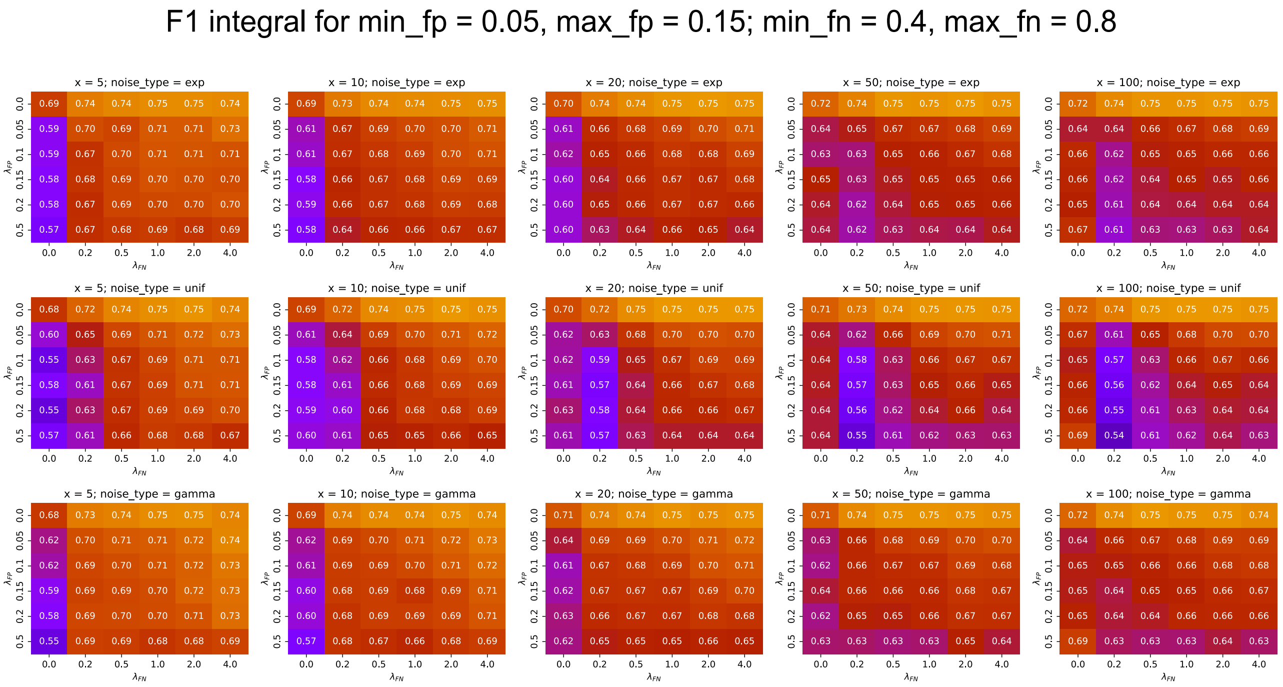


Figure 20: Normalised integral scores $\bar{S}_{F_1}$ across different noise augmentation regimes for the oxygen use phenotype.

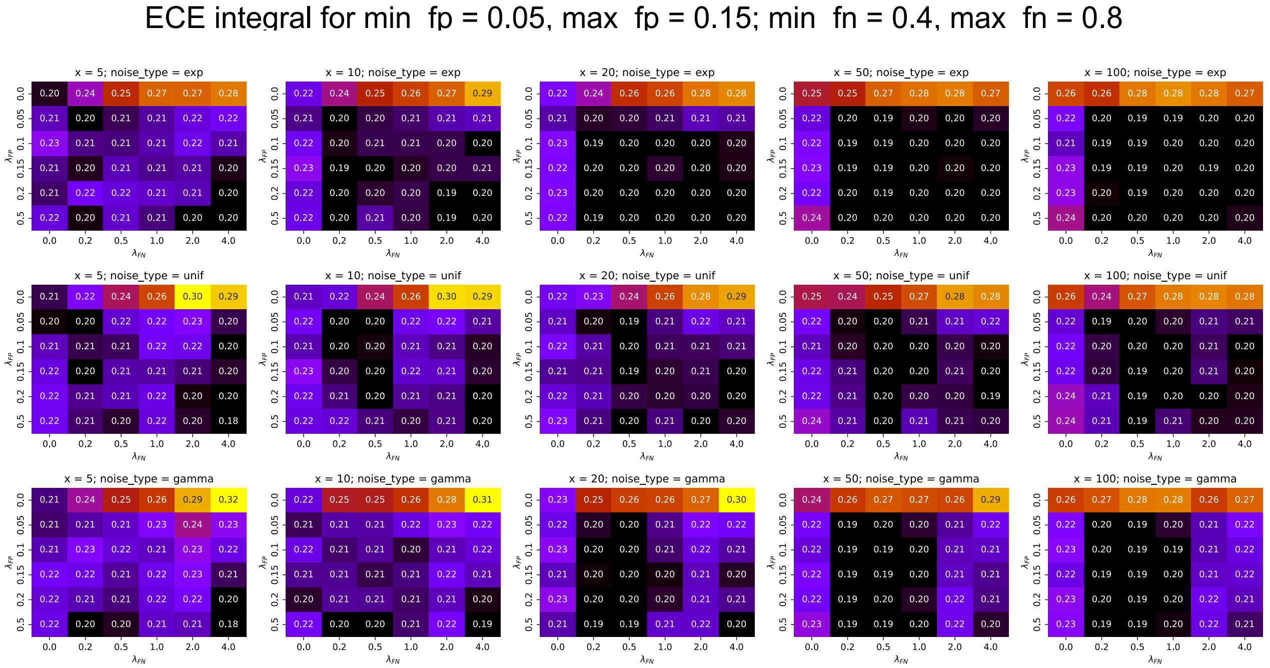


Figure 21: Normalised integral scores $\bar{S}_{\mathrm{ECE}}$ across different noise augmentation regimes for the oxygen use phenotype.

| | Binary class predict. | Predicted class proba. | Final regression predict. | 80% predict. interval | $\sqrt{\mathrm{Var}(\hat{y})}$ |
|---|---|---|---|---|---|
| Oxygen use | Anaerobe | 0.75 | - | - | - |
| Cell envelope | Diderm | 0.88 | - | - | - |
| Sporulation | Non-sporulating | 0.87 | - | - | - |
| OGT | Thermophile | 0.88 | 62°C | [46.2°C, 75.5°C] | 15.8°C |
| GC content | Low | 0.53 | 45 % | [37.7%, 54.1%] | 12.5% |

Table 7: LBCA phenotype predictions generated by models trained under the best-performing noise regimes identified in Table 6.

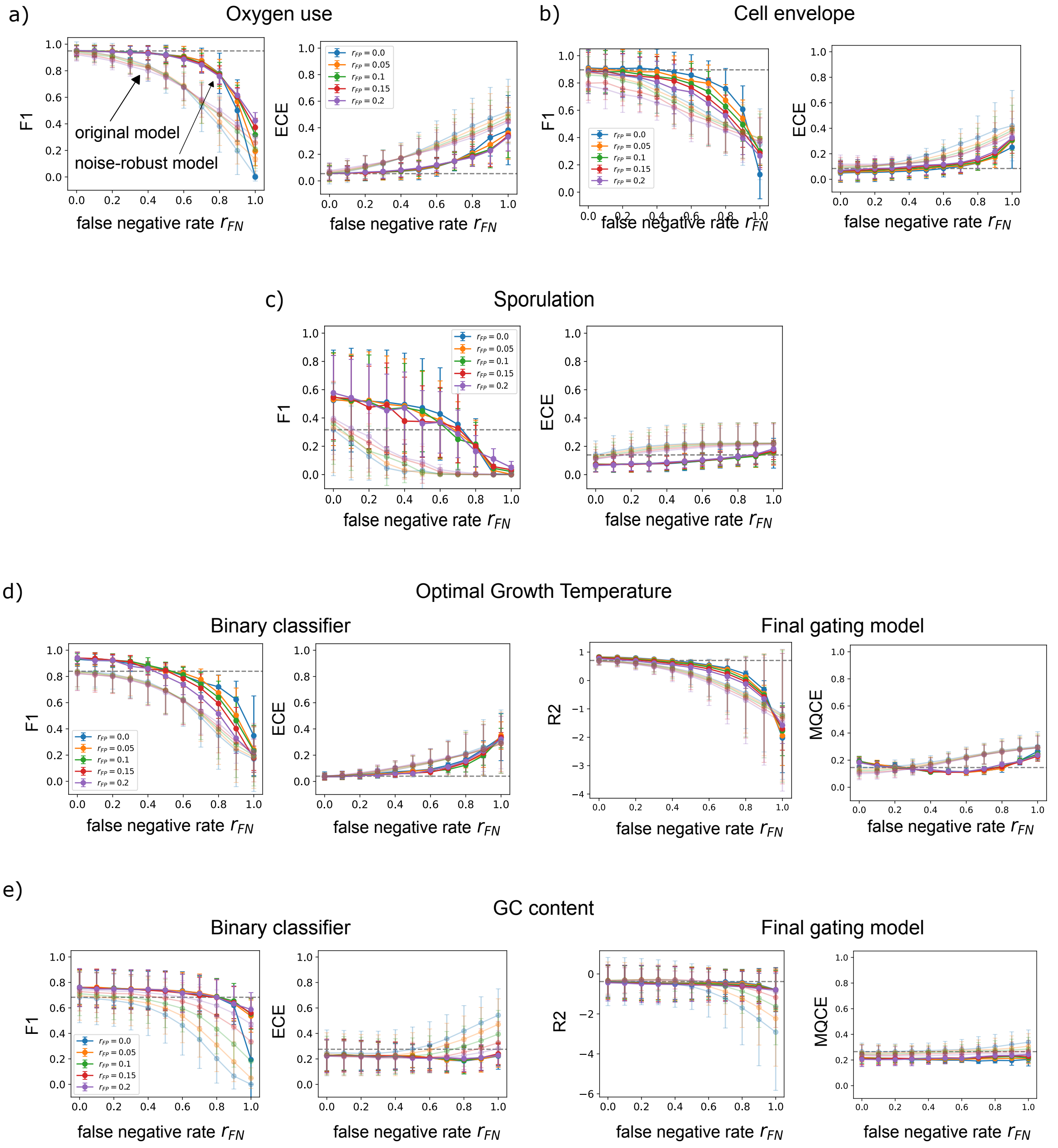


Figure 22: Performance comparison between the original models trained on noise-free datasets (semi-transparent lines) and the best noise-robust models trained on noise-augmented datasets (solid lines). Prediction accuracy is shown across different false-positive ($r_{FP}$) and false-negative ($r_{FN}$) noise rates applied to the test data. (a) Oxygen use. (b) Cell envelope. (c) Sporulation. (d) Optimal growth temperature. (e) GC content.

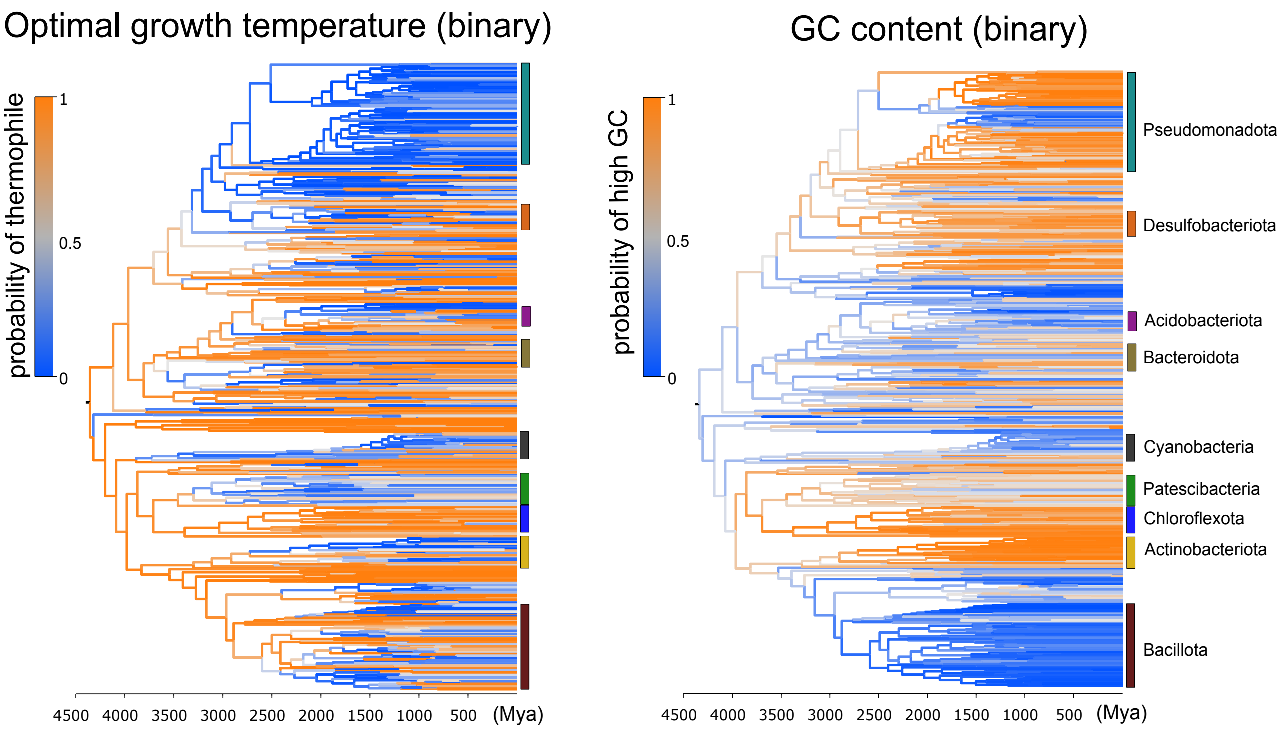


Figure 23: Ancestral classification of optimal growth temperature and GC content for the bacterial species tree from [**?**]. Branch colors represent the predicted class assigned by the binary gating classifier. (a) Optimal growth temperature (mesophile vs. thermophile). (b) GC content (high vs. low).
